# An inductive bias for generalization in mouse olfactory learning

**DOI:** 10.64898/2026.08.17.745288

**Authors:** Ningjing Xia, Venkatesh N. Murthy

**Affiliations:** Center for Brain Science and Department of Molecular C Cellular Biology, Harvard University, Cambridge, MA 02138, USA; Program in Neuroscience, Harvard Medical School, Boston, MA 02115, USA

## Abstract

Animals must generalize from limited experience, yet behavioral experiments in the laboratory setting rarely assess whether or how rapidly they generalize. This contrasts with machine learning systems, where generalization is considered a fundamental test of learning, and emphasizes performance evaluation with new in-distribution or out-of-distribution examples. Here, we used an olfactory categorization task to investigate rules of generalization versus memorization in mice. We trained mice to discriminate between two target odorants mixed with a variable number (0-13) of background odors. There are 32766 possible mixture stimuli to be classified, yet mice learn to generalize from as few as 8 unique mixtures. This generalization is not due to limited memory capacity: mice successfully learned to group the same set of mixtures when category labels were randomly shuffled. Analysis of individual variability revealed features in learning dynamics during training that predict performance in the generalization phase. A linear supervised learning algorithm could describe the generalization from few exemplars well, whereas nonlinear classifiers were necessary to explain memorization. Our experiments suggest that mice have an inductive bias towards generalization, consistent with a preference for simple rules, and will memorize only when forced to do so.

## Introduction

Neural networks are designed to learn^1,2^. Indeed, it seems that any complicated function or task can be learned using a complex enough model, given enough training data^3^. However, it remains uncertain how much the state-of-art models (including generative AIs) are still merely memorizing the training data^4–6^. Even if they are truly generalizing, the data required to train these models are orders of magnitude larger than what would be required by a human^7–9^.

Unlike artificial neural networks, animals, with their natural neural networks, are seen to require much less data to generalize^10–12^. This discrepancy has prompted the view^12^ that animal brains are not generic “blank-state” learners, but come equipped with powerful *inductive biases*, which are structured constraints and priors that narrow the space of hypotheses and thereby enable rapid generalization from limited data^10–12^. In parallel, work from machine learning also argues that end-to-end optimization over unstructured networks only triumph when the training resources are infinite, and that inductive biases are needed to both permit generalization in low data regime and enable models to think more like humans^11,13,14^.

These developments raise a central question for neuroscience: what are the inductive biases that animal brains implement, and how can we measure them behaviorally in a way that is commensurate with formal concepts such as generalization, sample complexity, and overfitting? Many studies have described the rapid learning and generalization of animals in different contexts^15–18^. However, these studies lack systematic control on the relevant inputs contributing to the learning process, and the learning process varies substantially between animals. Recent work on visual object recognition in rodents explicitly addresses the extent of generalization, but those studies still did not systematically investigate learning strategies and inductive biases^19–21^.

Contemporary systems neuroscience, with its focus on stable, well-controlled paradigms, has often favored repeated presentation of a fixed, small set of stimuli^22,23^. Under these conditions, high asymptotic accuracy is frequently taken as evidence that the animal has “learned the task,” even though the demands of the experiment may be met by memorizing a handful of contingencies. From the perspective of statistical learning, such conditions do not distinguish between an exemplar-based strategy and a rule-based one^24^, nor do they reveal how much diversity in experience is required before an animal will exploit its inductive biases to infer a more general solution.

Here, we seek to characterize inductive biases in mouse olfactory learning by explicitly contrasting memorization and generalization in a high-dimensional stimulus space. Rats and mice rely heavily on smell for survival, and their olfactory systems must parse high-dimensional, variable mixtures into behaviorally meaningful components^25–29^. We ask a quantitative question that can be posed equally to biological and artificial learners: how many distinct training examples are required before an animal abandons pure memorization and adopts a rule that supports broad generalization to novel stimuli?

To do so, we use an olfactory feature-detection task in which head-fixed mice detect the presence of specific target components within mixtures of many neutral odors, that act as distractors^25^. In our setup, the combinatorial stimulus space is enormous, allowing us to treat the tested mixtures as draws from an effectively continuous distribution rather than a finite list of items. Crucially, we manipulate only the *diversity* (defined as the number of unique examples) of the training set, while keeping the underlying categorization rule fixed, and we compare this regime to one in which the same stimuli are paired with arbitrary, non-rule-based reward associations. This design allows us to dissociate success driven by memorization of exemplars from success based on inference of a simple, compositional rule. Here, we use the term “rule” to refer to the rule of extracting the common component.

By measuring how performance on novel mixture stimuli depends on the richness of prior experience, and by demonstrating that animals can memorize arbitrary odor– reward assignments when required, we recover the inductive biases that guide mouse olfactory learning.

## Results

### Mice learn an odor mixture task from a small pool of training examples

To systematically study generalization in mice, we adapted an odor mixture discrimination task we developed previously^25,30^. In this task, the stimulus space of odor mixtures is combinatorial and large enough to test generalization with novel stimuli. The task design also allowed us to precisely and flexibly control the stimuli.

We delivered odor mixtures constructed from 16 odorants to head-restrained mice using custom-built olfactometers (**Figure 1a**, **Table S1** & **Figure S2**). Animals reported the presence of a Go target odor (ethyl tiglate) by licking a water spout or the presence of a NoGo target odor (ethyl propionate) by not licking (**Figure 1b**). All mixtures contained either the Go target or the NoGo target, mixed with various numbers of distractors (0 – 13). In correct Go trials mice were rewarded with a water drop. Incorrect trials had no punishment (**Figure 1b**). We focused on one pair of Go/NoGo targets because previous work showed no difference in performance when choosing different odors^25^. Any mixture contained 1 to 14 components, rendering a pool of 32766 possible stimuli (all odors are listed in **Methods**).

**Figure 1:**
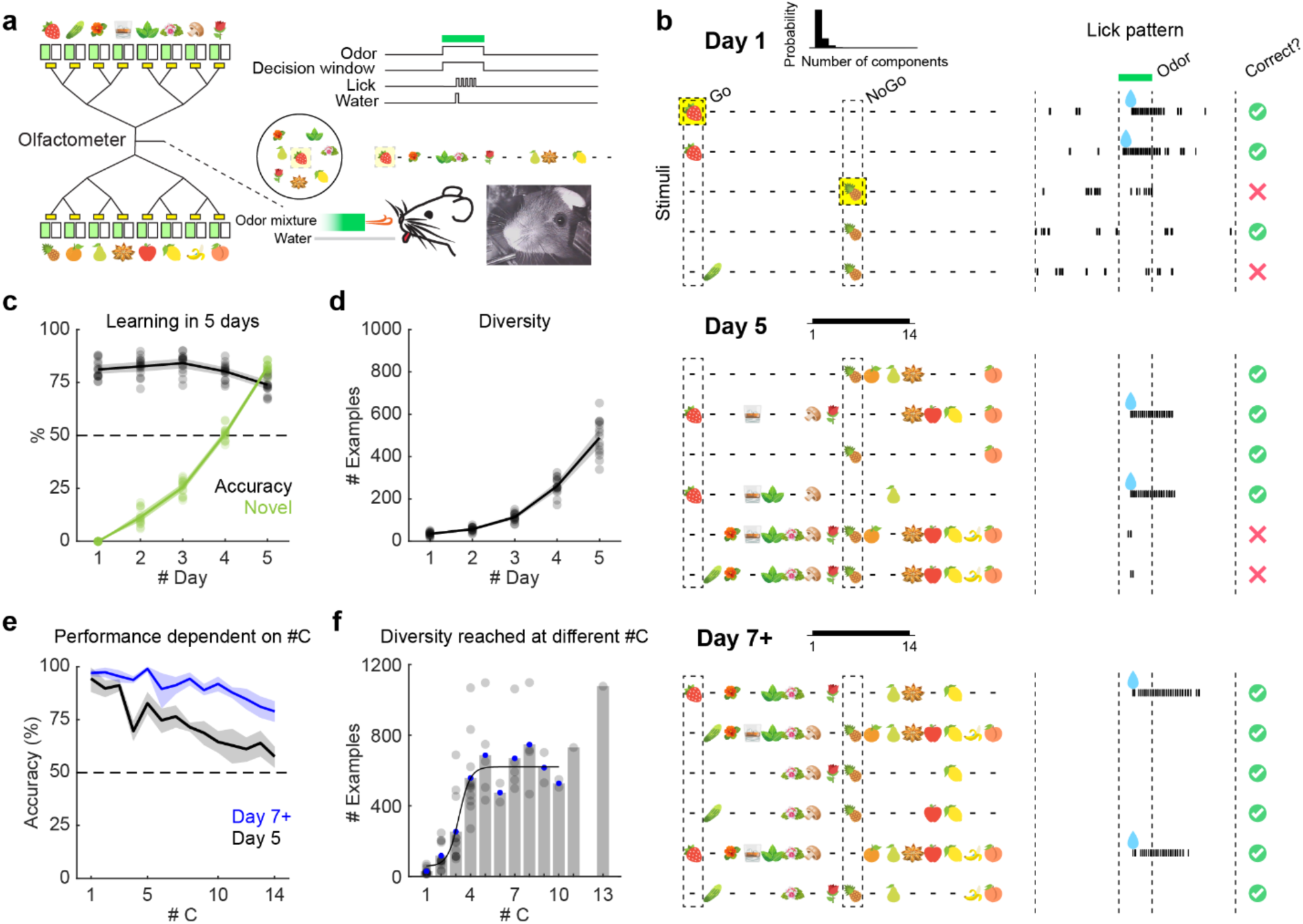
Mice display strong generalization in an olfactory mixture task. (**a**) Left: olfactometer, green: odor solvent, white: pure solvent, yellow: solenoid valve. Each of the 16 odorants is represented with an image symbol. Right top: task structure. Right bottom: illustration of odor and water delivery. (**b**) Example trials on Day 1,5, and 7 (or beyond 7). Upper panel black histogram: mixture distribution, x axis: number of components from 1 to 14, y axis: probability. Left: stimulus on each trial, middle: animal licking, right: correct or incorrect. Go target (strawberry) and NoGo target (pineapple) are marked with yellow squares on Day 1. Green bar on top represents odor presentation period. Day 1: most trials are single or low-#C stimuli. Day 5: #C is uniformly distributed between 1 and 14. Day 7+ (expert day): #C is still uniformly distributed between 1 and 14, but mice performance is near perfect, shown in (**e**). (**c- d**) Statistics of stimuli and mouse performance over 5 days of training (N=15 mice). Percentage of: correct trials (**c**, black), novel trials (**c**, green), and cumulative number of different stimuli encountered (**d**), on Day 1 to Day 5. (**e**) Different accuracy for different #C and performance improvement from the first day on uniform distribution of #C (Day 5, black) to the “expert day” (Day 7+, blue) across N=7 mice. (**f**) Different number of examples required to reach criterion performance for different #C trials, across 7-day training (N=12 mice). Gray bar: averaged # Examples required. Gray dots: each dot is data from one animal. Black line: fitted sigmoid curve. Blue dots: data used for fitting, the last two gray bars are omitted due to small N (#C = 11,13).

Mice learned the task in 5 days (accuracy > 75%) with a progressive curriculum (**Figure 1b-d**, N=15 mice; **Figure S3** & **Methods**). We show typical stimuli and animal responses on Day 1, 5, and beyond in **Figure 1b**. We collected data from four different experimental setups, each with a different olfactometer. Animals trained in one setup immediately performed well in another, indicating that they were responding to odors, rather than idiosyncratic cues specific to a given olfactometer (**Figure S4**).

Rapid learning implies good performance with limited time and limited experience. Across 5 days, mice would have encountered around 500 unique mixtures over roughly 1000 trials (**Figure 1d** & **Figure S3**). This is only ∼0.16% (500/32766) of the possible stimuli. Yet, mice responded with high accuracy to novel mixtures never presented before (**Figure 1c** & **Figure S3**, accuracy: 73.9 ± 1.10 %, percentage of novel stimuli: 82.1 ± 0.636 %, mean ± s.e.m., on Day 5, N = 15 animals). By the criteria in machine learning, this is strong generalization.

Adding more distractor odors increases the difficulty of detecting targets in proportion to the number of components (“#C”) in the mixtures since that leads to increasing probability of masking olfactory receptors that signal target odors^25,30^. Animals respond more accurately to low #C mixtures, and worse to high #C ones (**Figure 1e** & **Figure S3**). Additional training shifts the performance curve up, leading to a shallower dependence on #C (**Figure 1e** & **Figure S3**). Mice needed to encounter more examples to reach criterion performance (80%) for higher #C mixtures, but the number of examples reached a plateau at medium values of #C, reflecting generalization by mice (**Figure 1f**). That is, once mice have learned to detect target odors mixed with a threshold number of distractors, they can perform just as well for larger number of distractors.

In summary, we identified and quantified the remarkable generalization by mice in an olfactory feature detection task from only a small fraction of the total possible stimuli.

### Training-testing split experiments reveal generalization from few examples

How many distinct examples would animals need to encounter to reach a broad generalization on novel stimuli? To probe the dependence of the number of stimulus examples necessary for generalization, we adapted the standard train-test split from machine learning. Mice were trained on a fixed set of mixture stimuli. After training accuracy surpassed criterion, mice were tested on the full distribution of all possible mixtures to evaluate their generalization. We systematically varied the number of examples in the training set across different cohorts of mice (**Figure 2a-h**). We followed the machine learning design of training until asymptote, instead of controlling the same training time, as we want to directly probe the effect of training example diversity.

**Figure 2:**
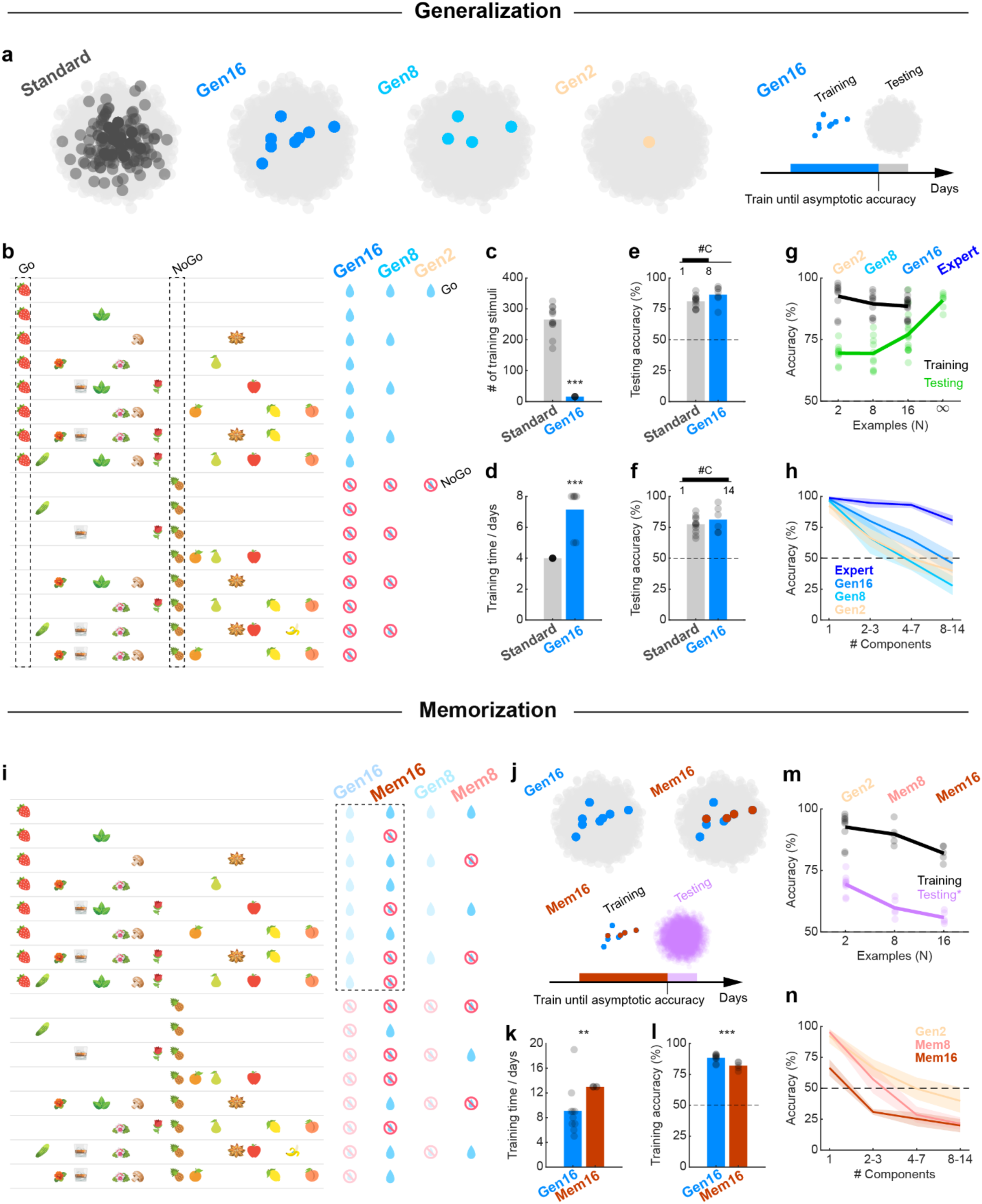
Generalization and memorization of the same mixtures reveal an inductive bias to generalize a simple rule in the mouse brain. (**a**) Illustration of the training-testing split design. Left, 4 dot clouds: training data are shown as 4 types of colored dots (Standard, Gen16, Gen8, Gen2, showing Go odors only) in the principal component space. The full distribution of testing data is shown with light gray dots. Right, time axis: animals are trained repeatedly on training data until criterion performance is reached, and then tested on the full distribution testing data. (**b**) The fixed stimuli presented to train different groups of mice in the generalization experiment. Each row represents a particular mixture, which is a combination selected out of 16 total odorants. The right most 3 columns indicate the reward contingencies for the stimuli in each of the 3 groups of mice. Gen16: 16 fixed stimuli were used as training data, N = 16 animals. Gen8: 8 stimuli, N = 10 animals. Gen2: only the two targets were used, N = 10 animals. (**c-f**) Training with a fixed set of 16 stimuli and testing on the full distribution (Gen16), compared to the unrestricted training curriculum (Standard)^25^. (**c**) Number of training stimuli encountered. ***p<0.001, bootstrapping over animals. (**d**) Training time. ***p<0.001, bootstrapping over animals. (**e**) Test day 1 (#C ∈ [1,8]). Standard: normal “training” day 5, trials satisfying #C ∈ [1,8] were taken for calculation. Gen16: test with all possible mixtures, #C uniform distribution from 1 to 8. (**f**) Test day 2 (#C ∈ [1,14]). Standard: normal “training” day 6. Gen16: test with all possible mixtures, #C uniform distribution from 1 to 14. (**g**) Generalization from 2, 8, 16 or many (“infinity”, or “Expert”) training examples, black: training accuracy, green: testing accuracy. Each dot is a different animal. Gen2: N = 10 animals. Gen8: N = 10 animals. Gen16: N = 16 animals. Expert (or “infinity”): N = 7 animals; Standard curriculum but longer training time (Day 7+) to illustrate expert performance. (**h**) Same in (**g**), showing different accuracy for different #C mixtures, only for the NoGo trials. (**i**) The exact same mixtures were used in the memorization experiments as in generalization experiments (**b**), but with shuffled categorization labels. Mem16: N = 5 animals. Mem8: N = 5 animals. Gen2: same data from Gen2 in (**g**), N = 10 animals. (**j**) Illustration of memorization experiment training scheme. The testing is same as in (**a-h**), but essentially new task for Mem16 and Mem8 animals. (**k**) Mem16 animals took longer to reach criterion performance than Gen16 in the training. **p<0.01, bootstrapping over animals. (**l**) Though both groups have passed criterion performance, the end training accuracy of Gen16 is higher than Mem16. ***p<0.001, bootstrapping over animals. (**m**) Black: training performance of memorizing 2, 8, 16 mixtures, lilac: testing on the mixture task rule with full distribution. Each dot is a different animal. Gen2: N = 10 animals. Mem8: N = 5 animals. Mem16: N = 5 animals. (**n**) Same in (**m**), showing different accuracy for different #C mixtures, only for the NoGo trials.

In the initial curriculum^25^ (referred to as the Standard curriculum hereafter), mice encountered >500 unique mixture stimuli during learning (**Figure 1**). In a radical departure, we next trained mice with only 16 examples (**Figure 2b**, **Gen16**, 2-test-day group, N = 7 animals). 8 of them were Go stimuli, and the other 8 were NoGo, with #C spanning from 1 to 8, covering different levels of mixture complexity (**Figure 2b**, **Gen16**). The same 16 stimuli were used for all mice in the Gen16 group. Mice were trained with the same overall curriculum as before, where low-#C mixtures occurred more frequently at first, and the probability gradually flattened to uniform distribution (**Methods** & **Figure S3**). Throughout training, animals encountered only these particular 16 mixtures in random sequence (**Figure 2c**). Remarkably, they performed as well in testing as those trained with an order of magnitude more examples (Standard curriculum group: 266 ± 13.2 stimuli from Day 1 to Day 4, mean ± s.e.m., Gen16 group: 16 stimuli throughout training, significantly different, ***p<0.001, bootstrapping, bootstrapping here and hence after is all done over individual animals, **Figure 2c**). On the first test day, #C ranged from 1 to 8, so mice were not challenged with high-#C mixtures that they never encountered before. On the second test day, #C spanned 1 to 14. Animals responded to test stimuli with above criterion accuracy on both test days (test day 1, Standard curriculum group: 81.1 ± 1.41 %, Gen16 group: 86.5 ± 2.70 %, mean ± s.e.m., p = 0.069 by bootstrapping, not significantly different, **Figure 2e**; test day 2, Standard curriculum group: 77.4 ± 1.91 %, Gen16 group: 81.3 ± 4.24 %, mean ± s.e.m., p = 0.11 by bootstrapping, not significantly different, **Figure 2f**). Interestingly, training on 16 examples required longer time to reach criterion performance than training with a larger number of examples (Standard curriculum group: 4 days, Gen16 group: 7.14 ± 0.55 days, mean ± s.e.m., significantly different, ***p<0.001, bootstrapping, **Figure 2d**).

Next, we reduced the size of the training set further to determine the smallest number of examples needed for animals to generalize in this task (**Figure 2b,g,h**). We trained different groups of mice on 2 (**Figure 2b, Gen2**, N = 10 animals) or 8 odor mixtures (**Figure 2b, Gen8**, N = 10 animals) until they reached criterion performance, and then tested their generalization. Animals are constantly learning, and we cannot freeze their internal weight-updating like in artificial neural network models. Therefore, we focused on just one testing day on the full distribution where #C spanned 1 to 14 (**Figure S5**). The Gen2 and Gen8 groups performed well in training, but did worse in testing compared to the Gen16 group (**Figure 2g**, **Gen16**, compiling 2- and 1-test-day groups, N = 16 animals). The two groups had similar average performance, while Gen8 animals displayed higher variation (**Figure S6**). In the Gen8 group, a few individuals exhibited well-above chance generalization in testing, whereas a few others performed worse than Gen2 animals. Test accuracy declined as mixture complexity increased – that is, mice trained with 2 or 8 odors performed progressively worse in trials with higher #C (**Figure 2h**).

Interestingly, mice continued to learn within the single session on the test day (in which many different stimuli were shown), such that the last 100 trials showed improved accuracy than the first 100 trials across groups (**Figure S7**). Under a strict definition of test stimuli, trials with the singles (Go target and NoGo target) or other mixtures that were presented during training should be excluded from the test accuracy calculation; doing so did not change the results significantly (**Figure S8**).

In summary, animals need a surprisingly low minimum diversity of examples in the training data to generalize. Experience with only 2 or 8 mixtures is not enough. However, training on as few as 16 examples renders good generalization on random samples drawn from the entire pool of ∼30,000 possibilities. We also note the high variability in test performance across individuals in some groups, especially Gen8, which we examine in later sections.

### Memorization experiments support the existence of an inductive bias

Animals trained with just a few examples can generalize well. This is counterintuitive because they could have memorized the training examples individually, and generalized poorly, since the memorization rule will rarely align with the generalization rule. Therefore, we asked if it’s possible for animals to remember the mixtures individually? To probe this, we asked whether animals could memorize the same specific (2, 8 or 16) mixtures with arbitrarily shuffled labels (**Figure 2i-n**). Our design was inspired by machine learning studies that define memorization capacity by measuring how many shuffled-label associations can be stored^6,31^.

Animals were trained with the exact same mixtures as in the generalization tasks, with a similar curriculum (**Methods**). However, the rewarding rules were changed, so that a simple rule of shared component no longer existed (**Figure 2i, Mem16**, N = 5 animals, **Mem8**, N = 5 animals). Therefore, mice could not systematically generalize, and were required to remember the mixtures individually. This is more evident looking at the Mem8 and Mem16 mixtures grouped by their categories (**Figure S9**). Mice reached high accuracy memorizing these shuffle-labeled mixtures (**Figure 2m & Figure S9**), but performance progressively dropped as more arbitrary category items were added (**Figure 2m**). Animals showed enough memory capacity to remember 8 or 16 mixtures, indicating that in the generalization tasks mice could have adopted a similar strategy to learn the Gen8 or Gen16 mixtures individually. However, the fact that Gen16 animals and a few individuals in the Gen8 group generalized from limited training examples suggested that the generalization is a result of animals’ inductive bias to extract a simple rule when a shared component defined a choice category. Mem16 mice took longer time than Gen16 ones to reach criterion performance (Gen16: 9.1 ± 1.3 days, Mem16: 13 days, mean ± s.e.m., significantly different, **p<0.01, bootstrapping, **Figure 2k**), and eventually reached a lower mean training accuracy (Gen16: 88.3 ± 1.0 %, Mem16: 82.0 ± 1.4 %, mean ± s.e.m., significantly different, ***p<0.001, bootstrapping, **Figure 2l**), suggesting that using the shared component rule is already advantageous with the 16-mixture training set.

If mice memorize the arbitrary associations between specific stimuli and outcomes, they should not perform well in the target-no target mixture generalization test since the memorization rule includes deliberate mismatch with the Go-NoGo rule (**Figure 2i,j**). Indeed, the Mem8 or Mem16 groups performed worse than mice just trained on the two targets, and performance dropped progressively as #C increased (**Figure 2m & 2n**). Both groups performed around chance level during the first 100 trials of testing session, which is a measurement that minimizes the effect of learning during testing (**Figure S8**). Learning a dissimilar stimulus-category rule even with a small number of samples interferes with learning the component membership rule.

These results indicate when mice experience a small set of stimuli, they display an inductive bias to quickly identify a simpler category rule instead of memorizing all possible members of the category, even if they possess the ability to do the latter.

### Training history features predict generalization versus memorization

Good performance in the testing phase for the Gen8 and Gen16 stimulus sets indicated that mice learned to generalize during training. We wondered whether there were identifiable features in the training history that would be indicators of generalization that already happened in the training phase. During training, animals learned the category labels of mixtures with different #C. If mice were using a generalization strategy, they would learn the easier – that is, low-#C –mixtures better, and find high-#C mixtures more difficult (**Figure 3a**, mouse “m47”). If mice were using a memorization strategy, high-#C mixtures would not necessarily be more difficult, because they might stand out with other salient features not related to the targets (**Figure 3a**, mouse “f54”). Any sign of disrupted order of learning is a sufficient, but not necessary condition of generalization.

**Figure 3:**
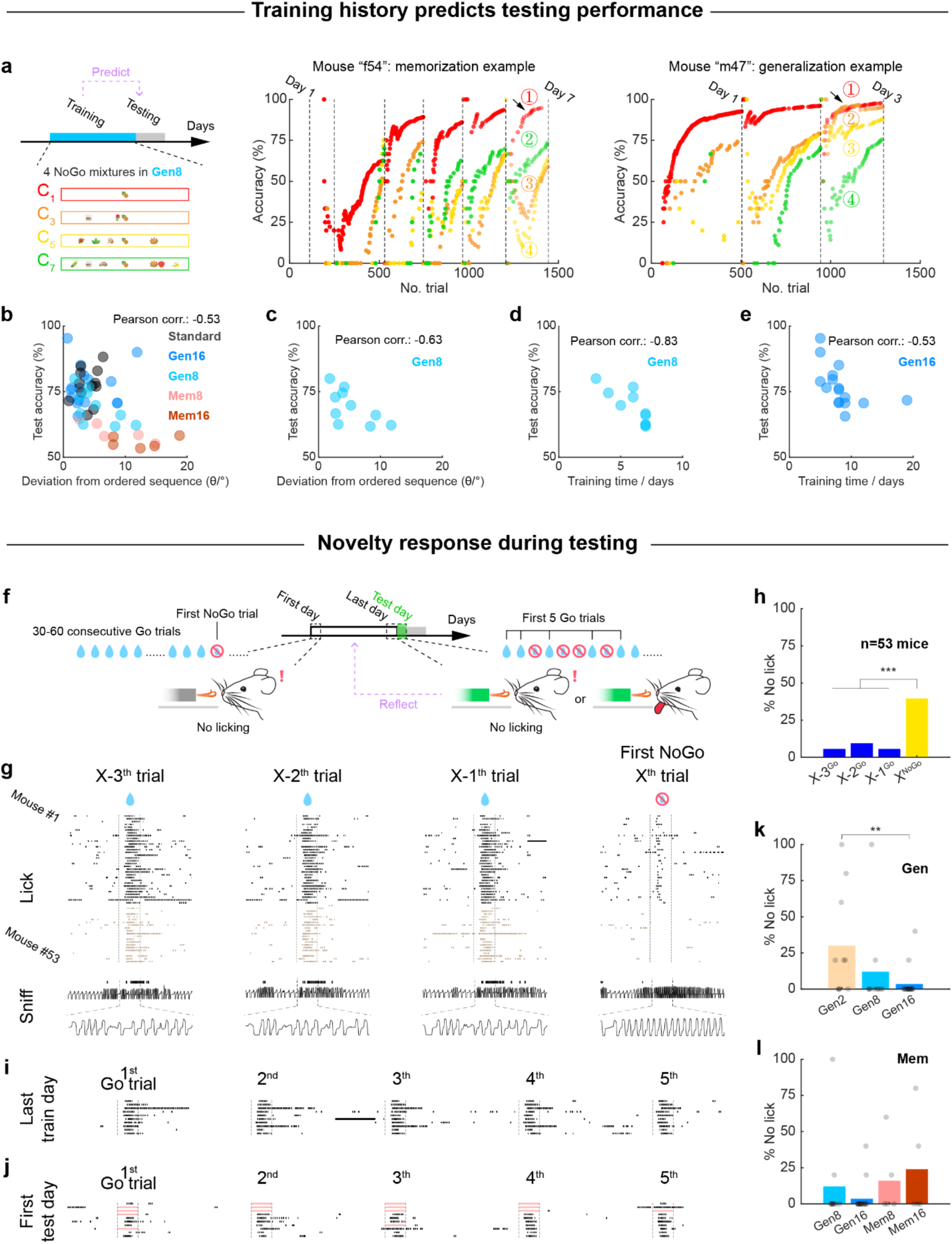
Behavioral metrics indicate different degrees of generalization. (**a-e**) Training statistics reflect different inductive biases and predict test accuracy of different individuals. (**a**) Illustration of the features in training histories that are extracted. Left: C_1_-C_7_: NoGo mixtures with #C = 1,3,5,7 in Figure 2a Gen8 group, colored in red, orange, yellow and green, respectively. Middle and right: example training histories (mouse “f54” and “m47”), only learning of NoGo mixtures is shown, dots: cumulative accuracy at each trial number, numbers in color: the order of training accuracies from highest to lowest, matching the C_1_-C_7_ natural sequence in mouse “m47” (1: red, 2: orange, 3: yellow, 4: green), and not matching that in mouse “f54” (1: red, 2: green, 3: orange, 4: yellow). (**b-e**): Correlation of test accuracy to (**b,c**) deviation from ordered sequence (θ), or (**d,e**) training time. Each dot is a different animal. (**b**) Pooling of all 5 groups of animals, N = 48 animals. (**c,d**) Gen8, N = 10 animals. (**e**) Gen16, N = 16 animals. (**f-l**) Mice making better generalization also display less novelty response to testing stimuli. (**f**) Illustration of novelty response (not licking) on the first training day and the beginning of the testing day. (**g-h**) Consistent display of novelty response by not licking to novel odor across N = 53 animals. (**g**) Pooling the response of 53 animals to their first encounter of the NoGo odor on the first day of training. X^th^ trial: the first NoGo trial, X is in the range of 30-60 and varies across different animals. X-3^th^, X-2^th^, X-1^th^ trial: the three consecutive Go trials before the first NoGo trial. Upper panel: licking, sorted by animals that licked (black) or not licked (brown) to the first NoGo trial, bottom panel: sniffing, example recording from 1 animal. (**h**) Quantification of (**g**): the percentage of animals not licking on the first NoGo trial (yellow) is significantly higher than the three preceding Go trials (blue). ***p<0.001, bootstrapping. (**i-l**) Mice trained with different training stimuli display different levels of novelty response at the start of testing session. (**i-j**) Pooling the response of the Gen2 group (N = 10 animals) to the first Go trials on the last training day (**i**) or the first testing day (**j**). Blue: lick signal, red: marking no lick during the entire odor presentation period. (**k-l**) Quantification across different groups for the percentage of no lick during the first 5 Go trials on the first testing day. (**k**) Animals are less surprised by test stimuli when exposed to more training examples. Gen16 group is significantly lower than Gen 2, **p<0.01, bootstrapping. (**l**) Comparing the groups trained on the same set of stimuli but with different rewarding rules. The memorization groups display increasing novelty response whereas the generalization groups display decreasing novelty response, as N increases from 8 to 16.

We used either a metric *θ* that we defined ourselves (**Figure 3b,c & Methods**), or the widely used statistical measure Kendall’s tau (**Figure S10**) to quantify how much mice deviated from the “ideal” accuracy patterns if mice were generalizing during training. Essentially, both *θ* and Kendall’s tau quantify how much the behavioral accuracy of mice to different-#C mixtures descend in order like in **Figure 1e** or **Figure 2h**, and both metrics generated similar results (**Figure 3b,c & Figure S10**). It’s worth noting that animals using a memorization strategy might also display accuracy patterns in descending order as #C increased, but a non-descending accuracy pattern most likely indicates memorization and not generalization. This is a sufficient but not necessary condition. We found a strong negative correlation between *θ* and test accuracy (Pearson correlation coefficient: -0.53, Spearman’s rank correlation coefficient: -0.45, **Figure 3b**), and similarly using Kendall’s tau. Using the *θ* metric in the training process, one can predict the animal’s performance in testing. *θ* was larger for the memorization groups, and smaller for the generalization groups. The Gen8 group, which is at the border of the transition from memorization to generalization, exhibited intermediate distribution of *θ*.

Interestingly, the *θ* or Kendall’s tau metrics also explained individual variability within a group (**Figure 3c & Figure S10**). The Gen8 group displayed a high within-group variability in the testing accuracy, despite showing a similar group average to the Gen2 group (**Figure S6**). Within the Gen8 group, the deviation index (*θ* or Kendall’s tau) during training (Pearson correlation coefficient: -0.63, Spearman’s rank correlation coefficient: -0.62, **Figure 3c & Supplement S10**), but not the training accuracy (**Supplement S11**), correlates strongly and negatively with the test performance. Different individuals in the group were trained with the same 8 examples and reached similar training success, but they appear to have used different underlying strategies to complete the training. The different values of deviation index in Gen8 showed that mice have different options to remember the small-size training examples: to memorize or to generalize, and therefore reflect the different inductive biases among different individuals. In Gen16, the uniformly small values of deviation index offer strong evidence that mice generalized with 16 examples, and explained why this group performed well in the testing.

Beside the deviation index, the length of training time to reach criterion training accuracy also reflects different strategies. In both Gen8 and Gen16 groups, the training time correlates strongly and negatively with test accuracy (Pearson correlation coefficient: -0.83, Spearman’s rank correlation coefficient: -0.86, **Figure 3d**; Pearson correlation coefficient: -0.53, Spearman’s rank correlation coefficient: -0.79, **Figure 3e**). We infer that if mice were memorizing, they would have to remember more items and therefore take a longer time to memorize each of the correlated mixtures. On the contrary, a generalization strategy, by extracting one common feature from all the examples, only requires one bit of memory and treats other features as variation of noise, and therefore is simpler and faster.

In summary, we found evidence in the training history that mice were indeed generalizing from a few examples. We also identified features in the training process that indicate a generalization versus memorization strategy, which can also explain within group individual variability, reflecting different inductive biases among different individuals.

### Novelty response to testing stimuli differentiates generalization versus memorization

The novelty response, or surprise at violation of expectation, can potentially distinguish a generalization versus memorization strategy. If mice were purely memorizing the training data, they would be surprised by the novel stimuli encountered in the testing phase. If mice were generalizing from the training data, they wouldn’t be surprised, as a new stimulus would be just another data point in the same distribution they expected.

When analyzing early trials on the first day of training, we noticed that some mice refrained from licking when new stimuli were introduced, likely because mice are cautious, neophobic animals (**Figure 3f**). In Go/NoGo tasks, once mice are trained to lick consistently, the challenge for them is to learn to refrain from licking to all stimuli. However, this indiscriminate licking did not occur at the start of training. On Day 1, animals were first presented with 30-60 consecutive Go trials. They learned to lick consistently, and were only exposed to Go odors in that period (**Figure 3f,g** & **Methods**). When the first NoGo trial was introduced, around 40% mice refrained from licking (fraction of not licking animals, three preceding Go trials: 6.92 ± 1.26 %, mean ± s.e.m., the first NoGo trial: 39.6%, significantly different, ***p<0.001, bootstrapping, **Figure 3h**). This was likely due to mice being neophobic and cautious about unfamiliar stimuli especially on the first day. In support of this idea, animals licked indiscriminately to all the trials at the beginning of Day 2, when all the stimuli would have been familiar (**Supplement S12**). Sniff recording showed significantly increased rapid sniffing to the first NoGo odor on Day 1, despite the animal choosing not to lick in that trial (**Figure 3g**). Since rapid sniffing is diagnostic of novelty^32,33^, we infer that the absence of licking to a stimulus is an indicator of novelty response to an unexpected stimulus.

We then used the absence of licking to ask whether and how much mice treated stimuli in the testing phase as novel or unexpected. At the beginning of the test session, many animals refrained from licking to both Go and NoGo trials (**Figure 3i,j**). We inferred that this could be due to mice being surprised by the variety of mixtures never presented before. We quantified the level of novelty response by the percentage of misses (no lick to Go trials) in the beginning period (first 5 Go trials) among all animals in each group (**Figure 3k,l** & **Methods**). We found decreasing level of novelty response as the number of unique training stimuli increased, which is consistent with animals exposed to fewer mixtures being more surprised (fraction of not licking, Gen2 group: 30 ± 11.6 %, Gen8 group: 12 ± 10.0 %, Gen16 group: 3.5 ± 2.6 %, mean ± s.e.m.; Gen2 and Gen8 group mean not significantly different, p = 0.24 by bootstrapping; Gen8 and Gen16 not significantly different, p = 0.48 by bootstrapping; Gen2 and Gen16 significantly different, **p<0.01, p = 0.0068 by bootstrapping, **Figure 3k**).

We compared the animals in the memorization experiment to those in the generalization experiment – both groups were trained with the exact same 8 or 16 mixtures but with different category labels. The memorization groups displayed higher level of novelty response when confronting new stimuli in the testing phase, consistent with them being surprised by stimuli that are not consistent with the rule they were trained with (fraction of not licking, Mem8 group: 16 ± 11.7 %, Mem16 group: 24 ± 16 %, mean ± s.e.m.; Gen8 and Mem8 group mean not significantly different, p = 0.82 by bootstrapping; Gen16 and Mem16 not significantly different, p = 0.17 by bootstrapping; Mem8 and Mem16 not significantly different, p = 0.70 by bootstrapping, **Figure 3l**). The Mem16 group also served as a control to show that despite being exposed to all the 16 possible odorants, mice still display high-level of novelty response because of the test rule contradicting the training rule.

In sum, we used an unexpected phenomenon of no licking to Go trials to quantify the level of novelty response, and found that animals exposed to fewer mixtures during training displayed qualitatively more novelty response.

### Linear readout from OSN representations recapitulates generalization from a few examples

Previous work showed that linear readout from the activity of olfactory sensory neurons (OSN), representing the neural inputs to the olfactory system, can solve the olfactory cocktail party task^30,34^. Here, we showed that a similar, simple linear readout can match behavior in this more challenging setting of generalization from a few examples.

We modeled the OSN activation pattern for each odorant as a random activation pattern matching the statistics of sparsity observed in experimental data^25,30^, and mixture representation as sigmoid saturation of linear sums of individual odorant responses (**Methods**). The modeled OSN representations had a scallop’s shape in a low dimensional embedding (**Figure 4a-c**). To mimic the task given to mice, we ask how many examples a model would need to reach the generalization of the same odor mixture discrimination task. We chose a simple model: a support vector machine (SVM), which uses a single trained weight vector of length N to perform categorization, where N is the number of OSN channels (**Methods**).

**Figure 4:**
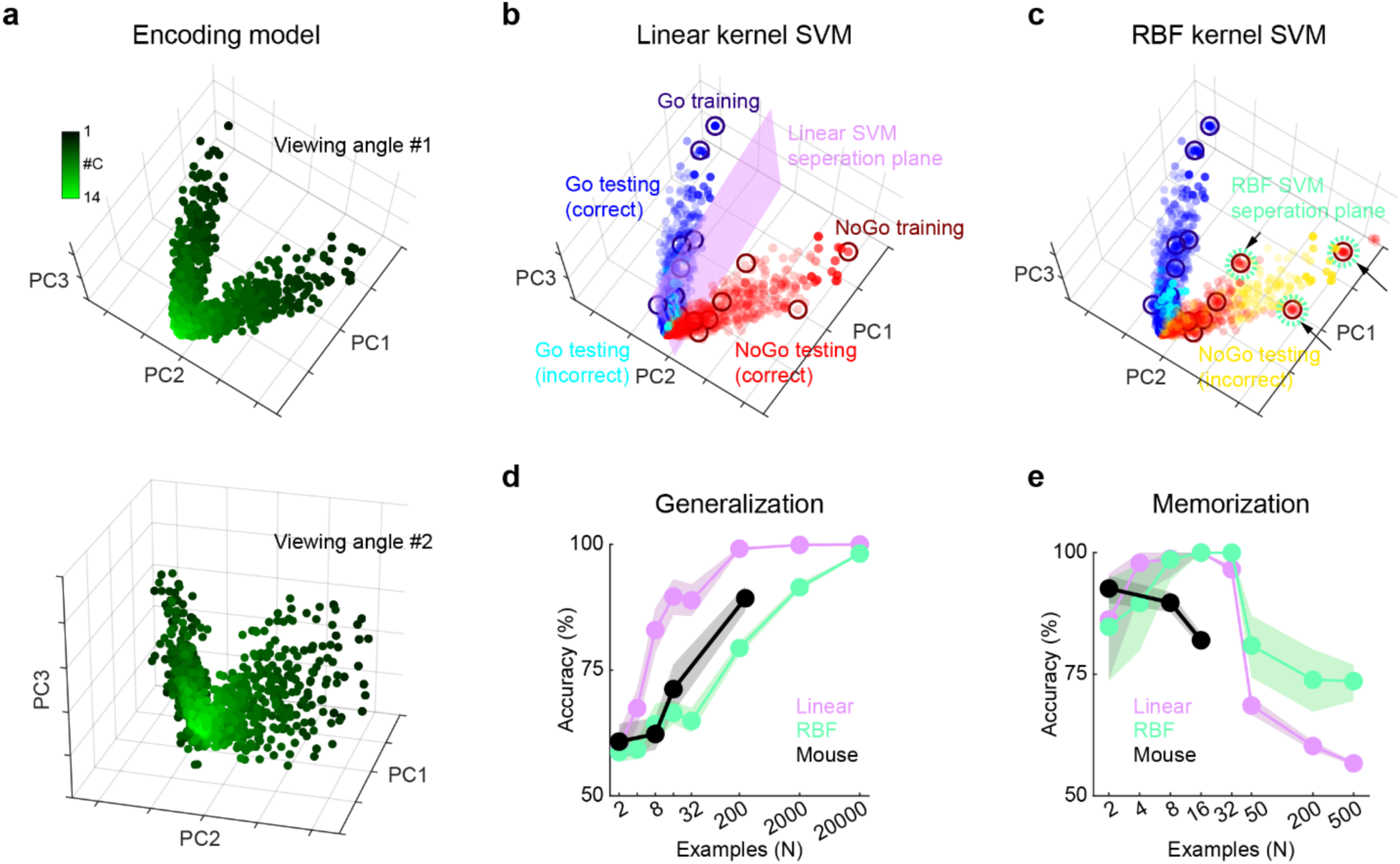
Performance of SVM models with linear or nonlinear kernels. (**a**) The encoding model of OSN activation patterns of odor mixtures in the PC space, spreading in a scallop-like shape. Dark to light green represents low to high #C. (**b-c**) Visualization of SVM with a linear or RBF kernel, trained on limited examples (dark, open circles) and tested on the full distribution (light, filled circles). (**b**) Linear kernel SVM. (**c**) Nonlinear kernel SVM. (**d**) Comparing the generalization curve of mouse, and two SVMs under different inductive biases. Mouse data also see in **Figure S8**. (**e**) Comparing the memorization curve of mouse, and two SVMs under different inductive biases.

An odor landscape contains multiple aspects of chemical information. One extreme way to process the information is to only care about the rule of the mixture task (extracting the common component) and throw out any other information, whereas another extreme is to be very accurate in remembering the particular odors and their relevant associations. We used SVMs with a linear kernel or a radial basis function (RBF) kernel to test which one mimics mouse behavior when limited training data are used. The linear kernel SVM will tend to discard detailed information about the stimulus and suggest simple hyperplanes for binary classification (**Figure 4b**), whereas the RBF kernel SVM will capture more details of each stimulus (**Figure 4c**). We tested SVMs with these two different kernels, or two different inductive biases, on how many different examples they require to generalize (**Figure 4d**). Interestingly, the linear kernel SVM performs very well with only a few examples (**Figure 4d**, lilac), cutting the separation plane right around the hinge of valves in the sensory input representation space (**Figure 4b**). On the contrary, the RBF (nonlinear) kernel SVM needs many more examples to improve accuracy on the testing data (**Figure 4d**, mint). Because the RBF kernel values locality, the separation plane carves out a sphere near a few closely clustered examples (**Figure 4c**). The generalization curve of mice is very close to the linear kernel SVM, suggesting that mice operate with an almost linear inductive bias (**Figure 4d**, black).

However, as the task demand changes from generalization to memorization, the strength of linear kernel SVM or RBF kernel SVM switches. In a different task resembling our memorization experiments, we ask the models to memorize increasing number of mixtures labeled randomly without any shortcut of membership rules (**Figure 4e** & **Methods**). Now the linear kernel SVM quickly collapses as the number of examples to remember becomes too large, but the RBF kernel SVM persists (**Figure 4e**), likely due to the locality advantage of RBF kernels.

In summary, while it is still a puzzle how the brain unites the synthetic and analytic aspects of odor mixtures, we showed that mice generalize at the almost linear inductive bias with a pair of simple SVM models.

## Discussion

Our results place mouse olfactory learning squarely within a modern, machine-learning– inspired framework to think about generalization. By explicitly separating training and test phases, and by exploiting the enormous combinatorial space of odor mixtures, we could ask a quantitative question usually reserved for artificial systems: how many examples does a learner need before it prefers to infer a rule rather than memorize exemplars? Three main conclusions emerge from our behavioral and modeling analyses. First, mice display striking generalization from very limited experience, often requiring on the order of only 10–20 mixtures to extrapolate correctly to tens of thousands of unseen stimuli. Second, the generalization exhibited by mice is not constrained by their memory capacity: animals can memorize arbitrary mappings for as many or more mixtures when a simple rule is not available. Third, the transition from memorization to generalization is graded and heterogeneous across individuals, and can be predicted from features of the learning history, suggesting that different mice deploy distinct inductive biases even in an identical task.

A central contribution of our study is the controlled demonstration that generalization in mice is nontrivial in a sense familiar from machine learning. In the standard curriculum, based on our initial study^25^, animals experience hundreds of distinct mixtures and, by the end of training, respond correctly on 80% of trials despite the fact that 80% of those stimuli are novel. Those experiments might suggest that a very rich experience is necessary for accurate performance. Perhaps, like over-parameterized deep networks, mice have effectively sampled the relevant stimulus subspace and can interpolate. The fixed-set experiments rule out this simple account. Animals trained only on 16 mixtures (which represents 0.05% of the stimulus space) can later perform almost as well as those with an order of magnitude more training exemplars. Reducing the training set further to 2 or 8 specific mixtures degrades test performance at the group level, but some individuals trained with 8 mixtures still generalize well. These results indicate that there is a threshold of *diversity* in experience beyond which mice are willing to invest in learning a compositional rule, and that this threshold varies across individuals.

Earlier work has indicated that rodents can extract abstract structure across successive odor discriminations, eventually solving new discriminations rapidly and with few errors^15,16,35–37^. These findings, analogous to the classic work of Harlow’s visual learning sets^38^, support learning of a rule, and imply that animals extract higher-order regularities – about which features are relevant, how stimuli are organized, and what kinds of rules are likely to govern reinforcement – well beyond mere storage of exemplars. Yet, even in these classical works, the size and diversity of the training sets were typically modest, and the relationship between memorization and generalization was not probed in a way that directly parallels modern concerns about overfitting in machine learning. Generalization has also been studied in the context of Pavlovian conditioning, but the training stimuli are limited^39,40^. Some recent studies have also started to apply the training-testing split in animal research^41,42^. Our work extends the above-mentioned studies in several ways. First, the dimensionality and diversity of our stimulus space is orders of magnitude larger: instead of dozens of well-controlled monomolecular odors, we operate in a combinatorial space of 32,766 mixtures. Second, we explicitly manipulate the *number of unique exemplars* seen during training while holding the underlying rule constant, providing a direct behavioral measure of sample complexity. Finally, by comparing the same mixture sets under structured (membership) versus random labeling, we can show that the switch from memorization to generalization is not simply due to capacity limits, but reflects an active preference for simpler explanatory hypotheses when they are compatible with experience.

The memorization experiments are particularly informative in this regard. When the membership rule is destroyed by shuffling labels, mice can still learn to classify 8 or 16 mixtures accurately, albeit with more training time, which is expected for arbitrary mappings. Thus, the bias to generalize of mice experiencing 8 or 16 stimuli under the structured rule cannot be attributed to insufficient memory resources. Instead, when exposed to a rich enough set of structured examples, mice *choose* to compress them into a feature-based rule, foregoing the need to store each item separately. When structure is removed, they revert to exemplar-based learning. This conditional deployment of generalization versus memorization supports the idea that animal brains embody strong task-specific priors^10,12,43,44^ – they will use them when the environment “invites” such structure, but are capable of falling back on rote memorization when no simple hypothesis fits.

Our analyses of the training histories provide an additional window into these internal strategies. The θ metric quantifies how closely an animal’s accuracy across mixtures of different complexity (#C) at the end of training matches an “ideal” pattern expected from rule-based learning, where mice showed high accuracy for simple, low-#C mixtures, and progressively lower accuracy for complex, high-#C mixtures, resembling the sequential learning of tasks with descending difficulties in machine learning^45^. Animals that eventually generalize well show small θ values: their learning curves unfold from easy to hard in a manner consistent with acquiring a common rule and then extending it. Animals with poor test performance, especially in the memorization conditions, display large θ values; they do not respect the ordering of complexity and treat some high-#C mixtures as no harder than simpler ones, as would be expected if each exemplar is learned independently. Importantly, θ predicts test performance even *within* a group trained on the same stimulus set, as in Gen8, whereas aggregate training accuracy does not. This dissociation indicates that traditional performance-based criteria (e.g. reaching 80% correct on the training set) may be blind to the underlying strategy. Two mice can score equally well during training yet differ dramatically in their ability to cope with novelty, because one has discovered a rule and the other has memorized a _list_^21,24,46^.

The dependence of test performance on training time reinforces this interpretation. In experiments using 8 or 16 stimuli, animals that took longer to reach criterion tended to generalize less well. A natural reading is that longer training reflects a heavier reliance on exemplar-based learning: it simply takes more time to accumulate quasi-independent associations to multiple specific mixtures than to infer a compact rule that applies broadly. In machine learning, similar trade-offs appear when comparing models with different inductive biases: architectures optimized for local interpolation can fit training data very well but generalize poorly, while those constrained to implement simpler hypotheses often generalize better and train faster^47^. That these relationships appear spontaneously across individual mice suggests that what we call “inductive bias” in artificial systems has a behavioral correlate in animal strategy.

Our novelty response measure provides a complementary, ethologically grounded probe of expectation. Mice are naturally neophobic and cautious about novel odors^48^, and early in training they sometimes refrain from licking when a new stimulus appears. We exploited this phenomenon by asking how often animals withheld licking to the first few Go trials during testing with the full distribution. Animals with impoverished training experience (Gen2 group) show much higher rates of such “misses” than those trained on richer sets (Gen16 group), consistent with the idea that the test distribution violates their expectations more severely. Moreover, memorization animals trained on 8 or 16 mixtures exhibit more novelty response at test than generalization animals exposed to the same mixtures but with a rule-consistent mapping. For these mice, the full-distribution mixtures are not simply “other examples from the same category,” but instead contradict a memorized pattern. Operationally, novelty response thus serves as a behavioral signature of whether the animal’s internal model treats new stimuli as in-distribution or out-of-distribution, a distinction that has become central in contemporary machine learning^49,50^.

The modeling results with linear versus RBF-kernel SVMs help to situate these behavioral phenomena in a computational landscape. When we encode mixtures as random, sparse OSN activation patterns with saturating nonlinearities, the resulting representation forms a “scallop-shaped” manifold in which mixtures cluster by target membership and complexity. A linear SVM trained on very few examples can find a decision boundary that slices this manifold near the hinge, misclassifying mostly high-#C mixtures, closely matching the error patterns of mice. By contrast, an RBF-kernel SVM, with its strong locality bias, tends to carve out local decision regions around each training point and performs poorly on unseen mixtures unless given many examples; even then, its error profile does not mirror the gradual degradation with #C seen in behavior. In other words, the pattern of successes and failures in mice is best captured by a model whose inductive bias is approximately linear at the level of OSN inputs.

At the same time, the memorization task reverses the comparison: now a flexible, highly local model like the RBF SVM better accommodates the random labels, while a linear classifier struggles as the number of memorized items increases. The brain, however, does not fully match either extreme. Mice can memorize 16 arbitrary mixtures, but not hundreds; they generalize efficiently from a few structured examples, but not with the brittle perfection of an idealized SVM. This intermediate profile suggests that olfactory circuits may combine linearly separable readouts optimized for feature detection, consistent with previous work showing feedforward decoding of mixtures from receptor inputs^30^, with more flexible associative mechanisms that support exemplar-like learning when necessary. Relating these algorithmic tendencies to specific circuit motifs in the mouse brain is a key challenge for future work.

Our findings resonate with, and extend, several broader themes in animal cognition. Studies in pigeons, bees, and corvids have demonstrated impressive categorization and concept learning, including transfer to novel exemplars and abstract relations such as “same–different”^51–53^. Much like Slotnick’s rats, these animals appear able to extract rules that transcend individual stimuli. However, the underlying stimulus spaces in those tasks are typically low-dimensional (e.g. simple shapes, colors, or numerosity), and the number of unique exemplars often limited. By placing mice in an explicitly high-dimensional, combinatorial space and carefully titrating stimulus diversity, we show that rule extraction is not restricted to simplified laboratory representations: it operates robustly in a domain that more closely approximates natural sensory complexity.

The work also has implications for how we interpret common behavioral assays in systems neuroscience. Many tasks used for probing decision circuits, value representations, or neural coding, such as two-alternative forced choices with a handful of stimuli, implicitly conflate memorization and generalization. Our results show that, even when animals achieve high and stable performance, their internal strategies can differ profoundly. Without explicit generalization tests or analyses of training dynamics, it is difficult to know whether a given neural correlate reflects rule learning, exemplar storage, or some mixture of both. Bringing machine-learning–style evaluation practices – train/test splits, out-of-distribution probes, and strategy-sensitive metrics – into routine use could sharpen the inferences we draw about neural computation.

In summary, by embedding a rich olfactory task within a machine-learning–inspired experimental design, we demonstrate that mice possess a robust inductive bias toward simple, feature-based rules, akin to an Occam’s razor operating in the olfactory system. This bias enables generalization from surprisingly few examples but is not obligatory: when the environment withholds simple structure, mice can resort to memorization. The interplay between these strategies, as revealed in learning dynamics, novelty responses, and model comparisons, provides a quantitative handle on the abstract notion of inductive bias in biological brains. The tendency of generalization from few examples probably depends on how stimuli are represented in the brain^54^. Future work combining this behavioral paradigm with large-scale neural recordings and targeted circuit perturbations will be essential for uncovering how these biases are implemented in the architecture of the olfactory system and how they might inspire more data-efficient artificial learners.

## Materials and Methods

### Animals

We used C57BL/6J mice (Jackson Lab ID: 000664) for most experiments. Occasionally (6 out of 15 mice in **Figure 1c,d**) Tbet-cre mice (Jackson Lab ID: 024507) were used and displayed no observable difference to C57BL/6J mice. Therefore, we pooled the data from all mice without specifying specific strains in the main text. We used both male and female mice, age ranging from 3 months to 2 years old. Sex or age did not show significant effects in our behavior paradigms. Specific genetic strain, sex, age or other information of all experiment animals can be found in the online sharing list of raw data. Animals are raised in a standard 12-hour light/dark cycle, along with other conditions complying with IACUC requirements.

### Surgeries

We performed surgeries to mount a custom-made titanium headplate on the animal’s skull for head restraint in later behavior experiments. Animals were anesthetized with ketamine/xylazine (100 and 10 mg/kg, respectively). We followed a standard procedure of opening the skin, gluing the headplate and further cementing (C&B Metabond). Post surgery, animals were administered with 1.3mg/ml extended-release buprenorphine (Ethiqa XR, Fidelis Pharmaceuticals #099114). Animals were monitored twice daily for at least four days after the surgery.

### Behavior

General design of the apparatus, the olfactometer, the odor set, and water delivery are described in previous work^25,30^. All the behavioral rigs were controlled using custom software written using LabView.

In each trial of the behavioral task, a 5-second pre-stimulus period was followed by a 2- second odor presentation, a 5-second post-stimulus period, and a variable inter-trial interval ranging from 5 to 10 seconds. Mice need to make decisions to lick during the 2- second odor presentation, with no additional imposed constraint on the timing. The first lick during this period marks the decision time. If it is a correct Go trial, water reward is immediately given after the first lick.

After at least 1 week of recovery from surgery, mice were subjected to water restriction for at least 7 days before starting behavior training. Every day mice were weighed and monitored. Around 1-2 ml of water is given daily to keep the weight loss stable and above 85%. Habituation to the experimenter’s hand also happened in this period, such that mice could comfortably groom while on the hands. For some mice, we performed a habituation to head restriction for 1 or 2 days; other mice did not go over a head restriction habituation but could adapt quickly on the first day of training. After the weight loss was stable, we started the odor training. Mice were head restrained and presented with an odor tube and a waterspout. We did not perform the typical “lick training”, where no odor was given and mice were trained to lick consistently from the waterspout. Instead, mice always received water rewards paired with odor presentation. Different groups of mice were trained with different sets of mixture stimuli. A special Day 1 training is shared across all mice, following the protocol and advice from Dr. Gonzalo Otazu^55^. On Day 1, mice encountered odor and water reward for the first time in the apparatus. We typically gave 30 trials of Go trials with free water (water would be given 700 ms after odor onset, even when animals have not licked; if animals licked before 700 ms after odor onset, they received water reward immediately, after the first lick), 30 trials of Go trials without free water, and then started introducing NoGo trials which are randomly mixed with Go trials. On Day 1, most mice learned to lick to the Go odors after the consecutive Go trials, and started learning the discriminated association. Some mice did not lick too much on Day 1. For these animals we repeated the procedure on Day 2 or Day 3 until they started reliably licking after the odor presentation. Except for the first chunk of Go trials on Day 1, the ratio of Go versus NoGo trials was kept at 1:1 throughout the training. Except for the first chunk of Go trials on Day 1, we also had an additional constraint where we did not let more than 3 (≥4) trials to be the same type (all Go or all NoGo) to maintain the animals’ attention.

#### Standard curriculum

As in previous work^25,30^, we controlled the difficulty of odor mixtures with the statistics of their number of components (*x*):

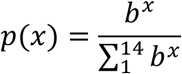

where *p*(*x*) is the probability of mixtures with *x* components. In this default curriculum, we draw stimuli from the entire distribution under different parameters of *b* without restrictions like in the following experiments. Typically, on Day 1 and 2, we used an easy level of difficulty with *b* = 0.25. On Day 3, *b* = 0.5. On Day 4, *b* = 0.75. From Day 5, we used the highest difficulty of *b* = 1, meaning that *x* is drawn from a uniform distribution from 1 to 14. Some mice were kept on this highest difficulty for some extended number of days.

#### Generalization experiments

Animals in the Gen16, Gen8, Gen2 groups only encountered a fixed set of mixtures during training. The Gen2 group received 4 days of training on just the Go and NoGo targets (single odors). The Gen8 and Gen16 groups were trained with 8 or 16 mixtures, respectively, and the statistics of these mixtures follow the Standard curriculum (**Supplementary Figure S3**). The probability of occurrence (*p*(*x*)) of a mixture with *x* components is calculated as follows:

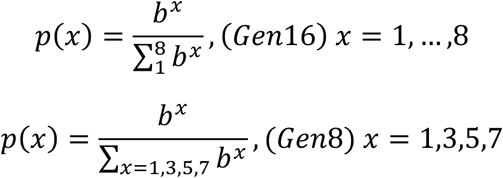

Similarly, we gradually increased the difficulty by changing *b*. Animals are considered to have completed training when their performance on the uniform distribution (*b* = 1) is above ∼80%. Then we tested their performance on the full distribution, with the uniform distribution, like the most difficult level in the Standard curriculum.

#### Memorization experiments

These were similar to the Generalization experiments (**Supplementary Figure S9**), except that the Mem16 group was trained with more sequential, incremental training statistics, where animals were first taught one pair of mixture associations, and then more, until they reached above ∼80% accuracy for all 16 mixture associations.

#### Calculation of the *θ* metric

We first calculated the “observed vector”, [C_1_,C_2-3_,C_4-5_,C_6-8_], where C_i-j_ is the accuracy of NoGo trials for mixtures with #C ranging from *i* to *j*. The lower bound of 8 was chosen because in Gen8 and Mem8 experiments the #C was capped by 8. Specifically, for the five groups to perform the calculations the observed vectors are: Standard curriculum, Gen16 and Mem16 groups: [C_1_,C_2-3_,C_4-5_,C_6-8_], Gen8 and Mem8 groups: [C_1_,C_3_,C_5_,C_7_].

Next, we calculated the “ideal vector”, where we kept the ratio fixed between the supposedly highest (C_1_) and lowest (C_6-8_) elements, and let the two middle elements (C_2-3_,C_4-5_) lie in sequence of gradually decreasing:

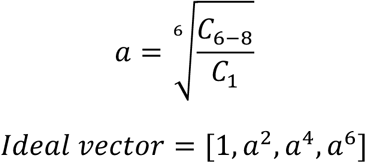

Finally, the *θ* metric is calculated by the angle between the observed vector and the ideal vector:

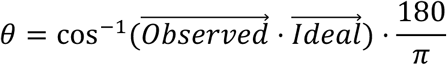

#### Calculation of the novelty response metric

For each animal, we calculated the percentage of misses (no lick to Go trials) in the first five Go trials on the first testing day. The group novelty response metric is the average of this number across all animals in the group.

#### Sniff recording

Surgery and sniff recording during behavior is same as previously shown^37^.

### Modeling

#### The encoding model of OSN representations

Based on previous work^30^, we construct the OSN representations of mixtures by applying a sigmoid nonlinear saturation to the linear sums of single odorants.

We generated the patterns of OSN activation for single odorants as follows:

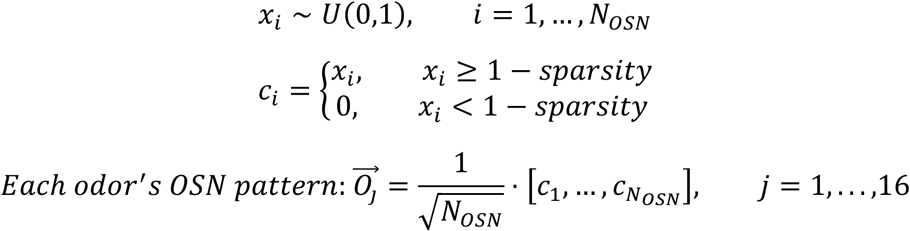

U(0,1): uniform distribution between 0 and 1, N_OSN_ = 100, sparsity = 0.4.

We generated the patterns of OSN activation for mixtures as follows:

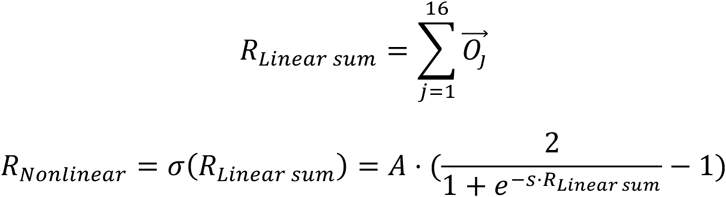

#### Linear and RBF kernel SVM

We use the Matlab function *fitcsvm* to fit SVM models, with either linear or RBF kernels (attribute “KernelFunction”).

#### Generalization experiments

We constructed training data like in our mouse experiments. To be specific, for # Example = 2,8,16, we used the exact same set of stimuli as in mice experiments, for # Example = 4,32, we designed the stimuli under similar principle, and for # Example = 200,2000,20000 as well as the testing stimuli (N_Testing_ = 2000), we constructed the stimuli similar to the last-stage curriculum of Standard curriculum, where #C is uniformly distributed between 1 and 14.

#### Memorization experiments

The training data was the same as in the generalization experiments, except that we tested smaller # Example up to 1000, and that the category labeling was random and not conforming to any membership rule. The testing data was generated by perturbing the training data in its vicinity by a small magnitude (0.1).

## Data analysis and Statistics

All the codes to analyze data or generate figures were in Matlab. Because most data were skewed and bounded between 0 and 1 (for example, accuracies or ratios), for most statistical testing, we used bootstrapping, over each individual animal as the independent unit. In suitable circumstances, we used t-tests.

## Data sharing

Raw data and codes will be shared online post publication.

## Acknowledgements

We would like to thank Ed Soucy and Yuwei Li for their technical support. We thank Gonzalo H. Otazu, Honggoo Chae, Jacob Zavatone-Veth and Naoshige Uchida for helpful discussions. We thank Samuel Gershman, Dan Rokni, Vikrant Kapoor, Farhad Pashakhanloo, Siddharth Jayakumar, Jacob Zavatone-Veth and other members of the Murthy lab for valuable feedback. Research in V.N.M.’s lab related to this manuscript was supported by grants from the NIH (RF1NS128865) and NTT Research (A47994). This work has been made possible in part by a gift from the Chan Zuckerberg Initiative Foundation to establish the Kempner Institute for the Study of Natural and Artificial Intelligence at Harvard University. N.X. was partially supported by the Mind/Brain/Behavior Interfaculty Initiative at Harvard University.

## Author Contributions

N.X. and V.N.M. conceived the project. V.N.M. supervised the project. N.X. designed the experiments, built the experimental setup, collected and analyzed the data reported in this paper, with input from V.N.M. N.X. implemented the model. N.X. and V.N.M. wrote the manuscript.

## Supplemental table and figures

**Figure S2:**
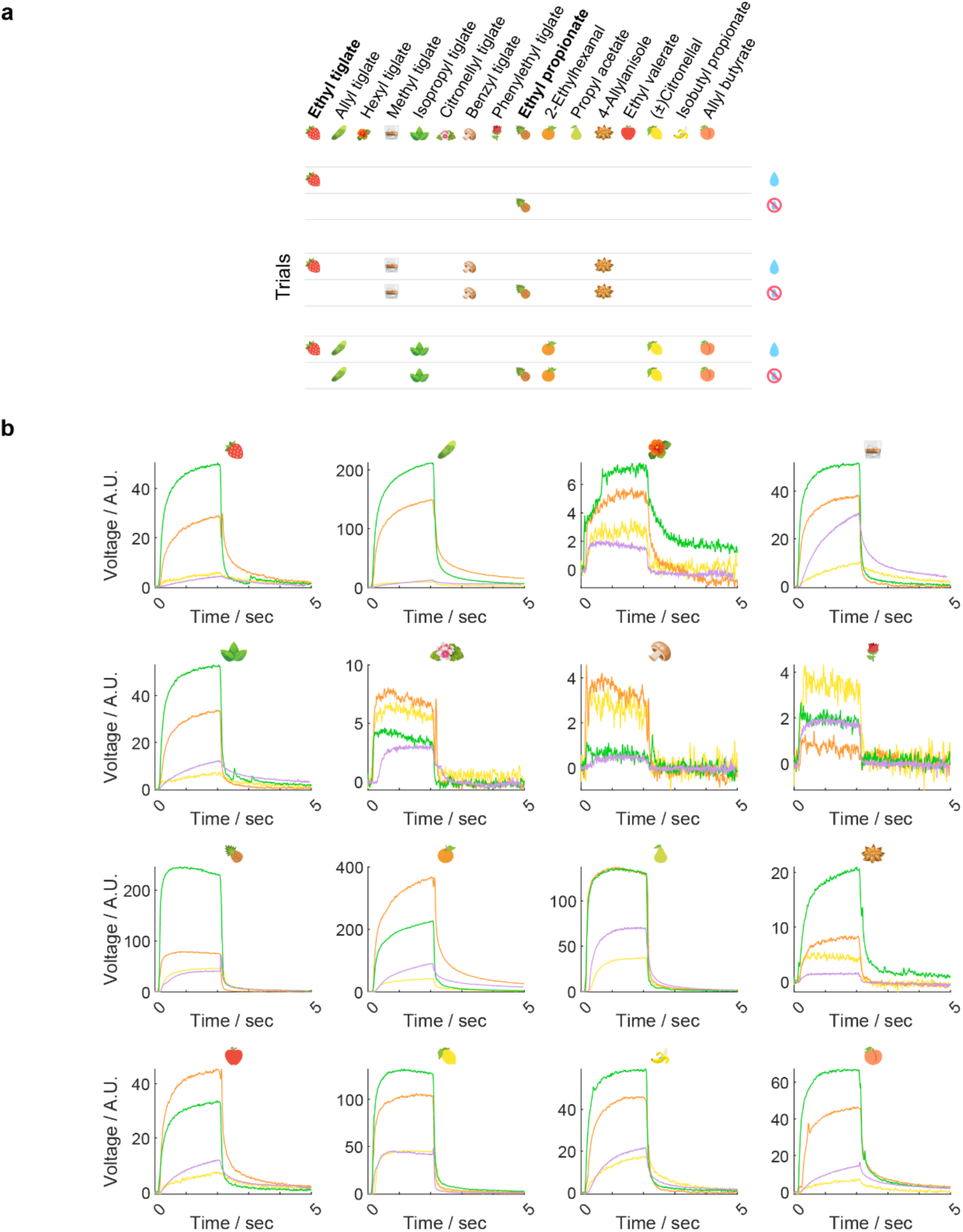
Examples of the task rule and PID of four setups. (**a**) Examples of task rule to categorize mixtures. (**b**) PID data testing the release of the single odorants in four olfactometers, each in a different color (yellow, orange, green, purple).

**Figure S3:**
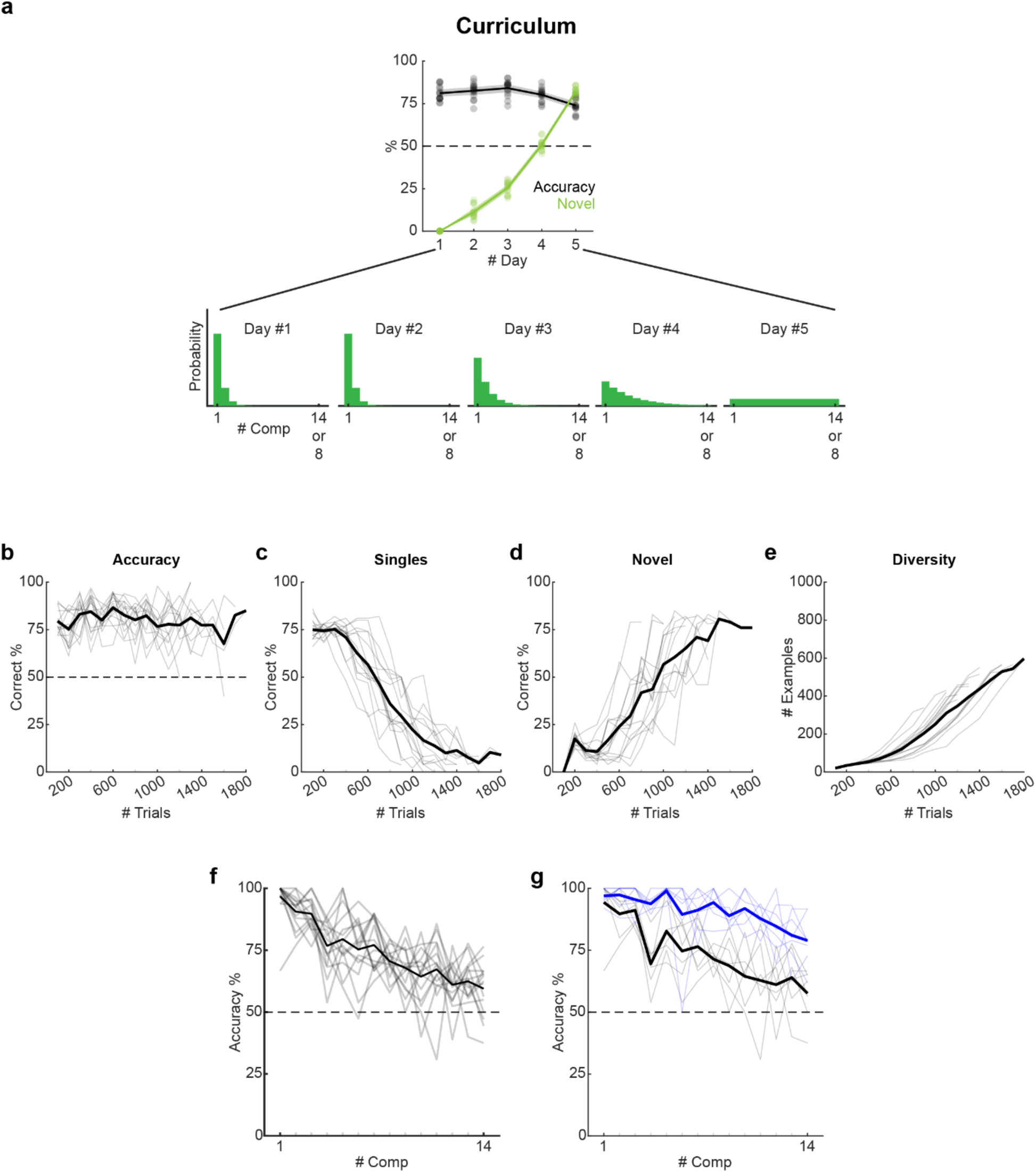
Training curriculum in 5 days. (**a**) The typical training curriculum used in a 5-day training. The probability of stimuli presented starts with being heavily skewed to easy, low-#C mixtures and gradually shifts to a uniform distribution. Depending on different experiments, the number of components is bounded by 14 or 8. (**b-e**) Learning statistics binned not by day, but by trials (100-trial bin), in N = 15 animals. (**b**) Like Figure 1c, black lines, showing individual animal’s trace. (**d**) Like Figure 1c, green lines, showing individual animal’s trace. (**e**) Like Figure 1d, showing individual animal’s trace. (**f**) Like Figure 1e, black lines, showing individual animal’s trace in N = 15 animals. (**g**) Like Figure 1e, black and blue lines, showing individual animal’s trace in N = 7 animals.

**Figure S4.**
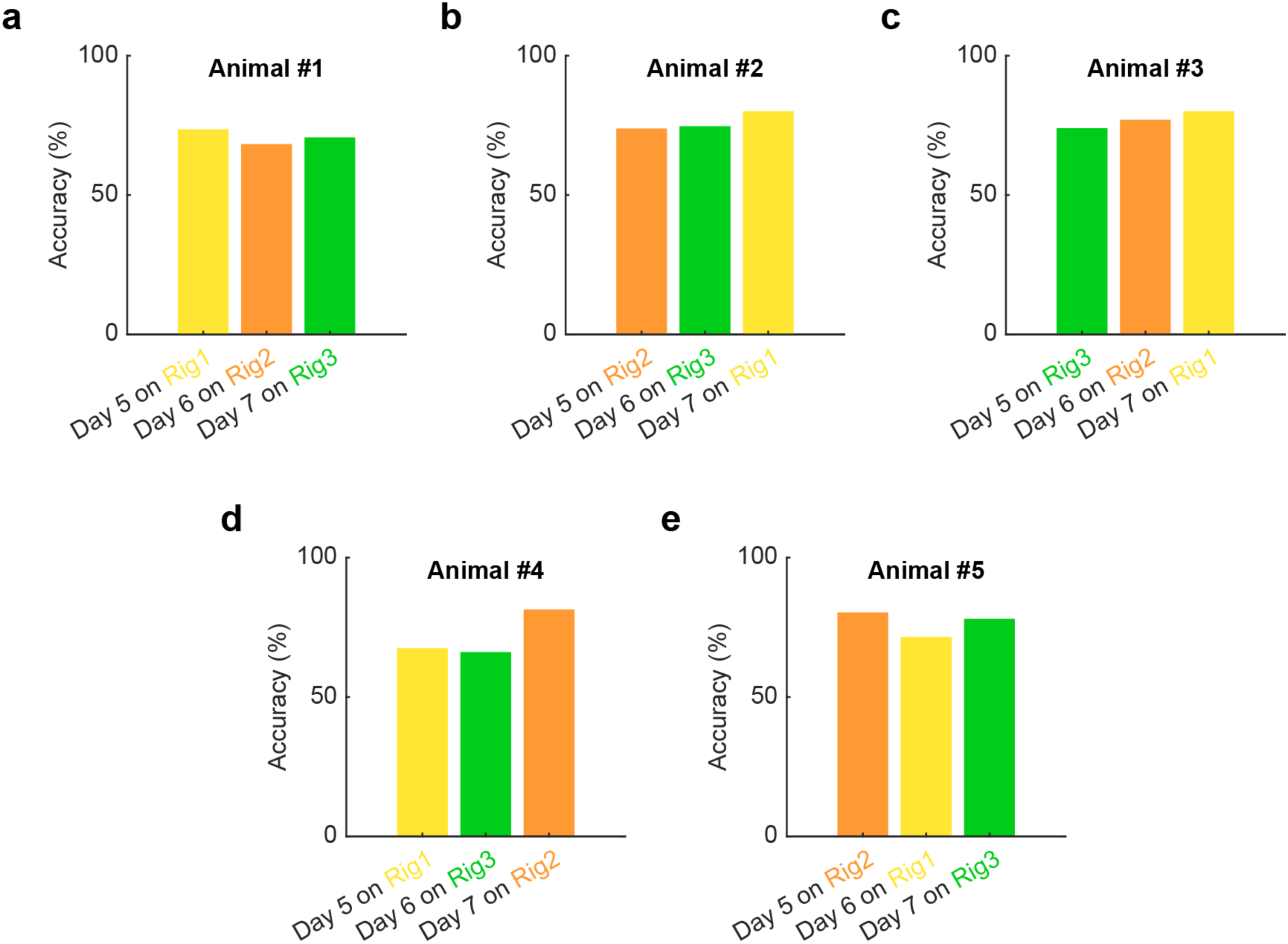
Animals trained on one setup can immediately perform in another. (**a**) An animal trained on Rig1 (yellow) from Day #1 to Day #5 reached the uniform distribution curriculum on Day #5. On Day #6 the animal was moved to Rig2 (orange) and trained on the uniform distribution. On Day #7 the animal was moved again to Rig3 (green) and still trained on the uniform distribution. The performance was similar across the three different rigs. (**b-d**) Like (**a**), each showing a different mouse trained on different sequence of the three rigs.

**Figure S5:**
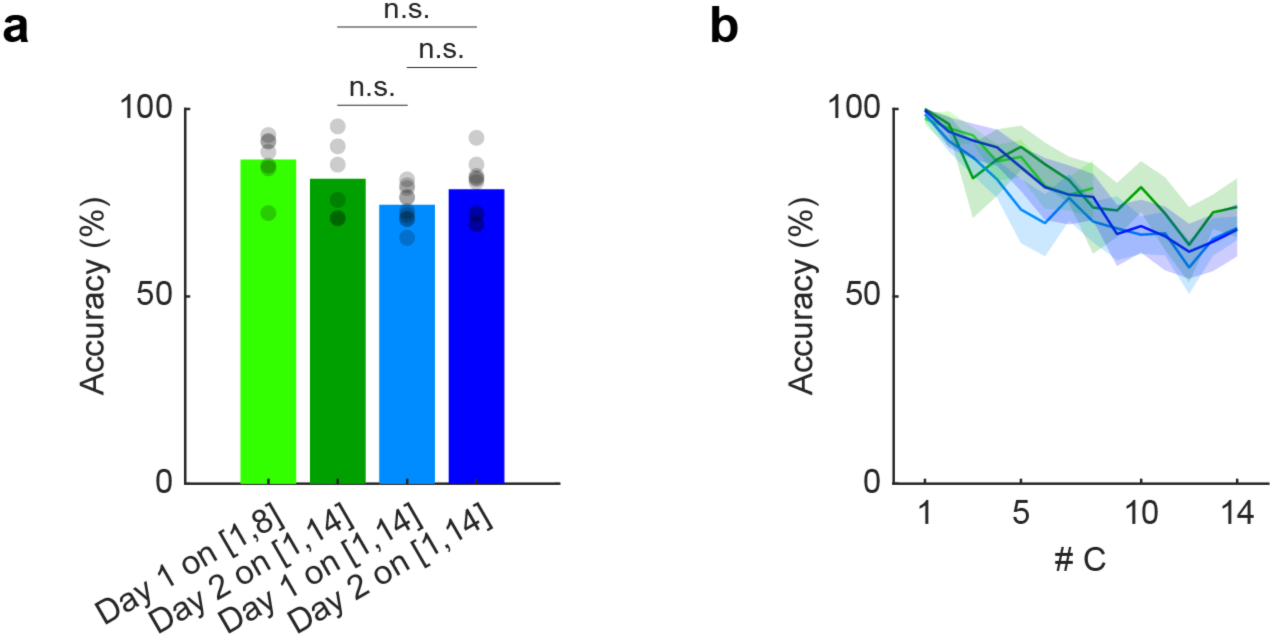
No significant difference between data that were pooled from two test days. (**a**) Data from two batch of animals showed no significant difference and were pooled together. Green (light and dark): N = 7 animals trained on Gen16 and tested with #C∈[1,8] on test day #1 (light green), and with #C∈[1,14] on test day #2 (dark green). Blue (light and dark): N = 10 animals trained on Gen16 and tested on #C∈[1,14] on both test day #1 (light blue) and test day #2 (dark blue). Dark green bar to light blue bar: p = 0.086 by bootstrapping. Dark green bar to dark blue bar: p = 0.54 by bootstrapping. Light blue bar to dark blue bar: p = 0.13 by bootstrapping. (**b**) Showing the different accuracies to different #C.

**Figure S6.**
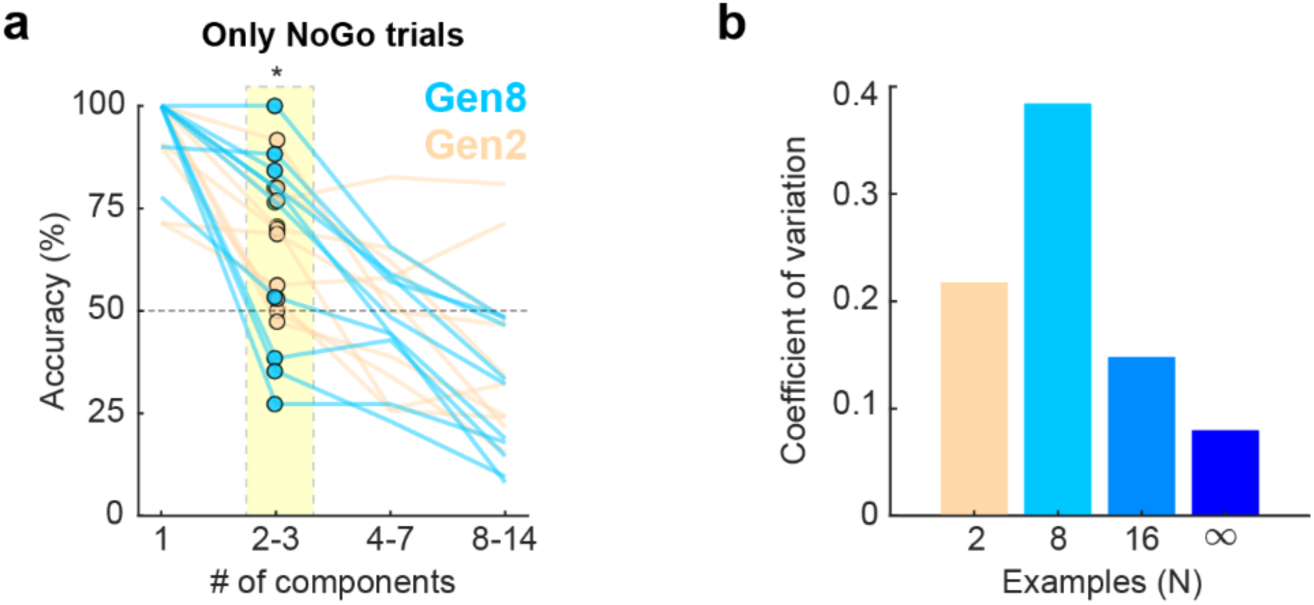
Gen8 displayed high within-group variability. (**a**) Different accuracy for different groups of #C, restricted to only NoGo trials. Blue: Gen8, peach: Gen2, each trace: individual animal, dots in highlighted square: the Gen8 group and Gen2 group display significantly higher variance despite similar mean in the low #C (2-3) trials, *p<0.05, bootstrapping. (**b**) Quantification of data in (**a**) by coefficient of variation, for #C = 2-3, NoGo trials.

**Figure S7:**
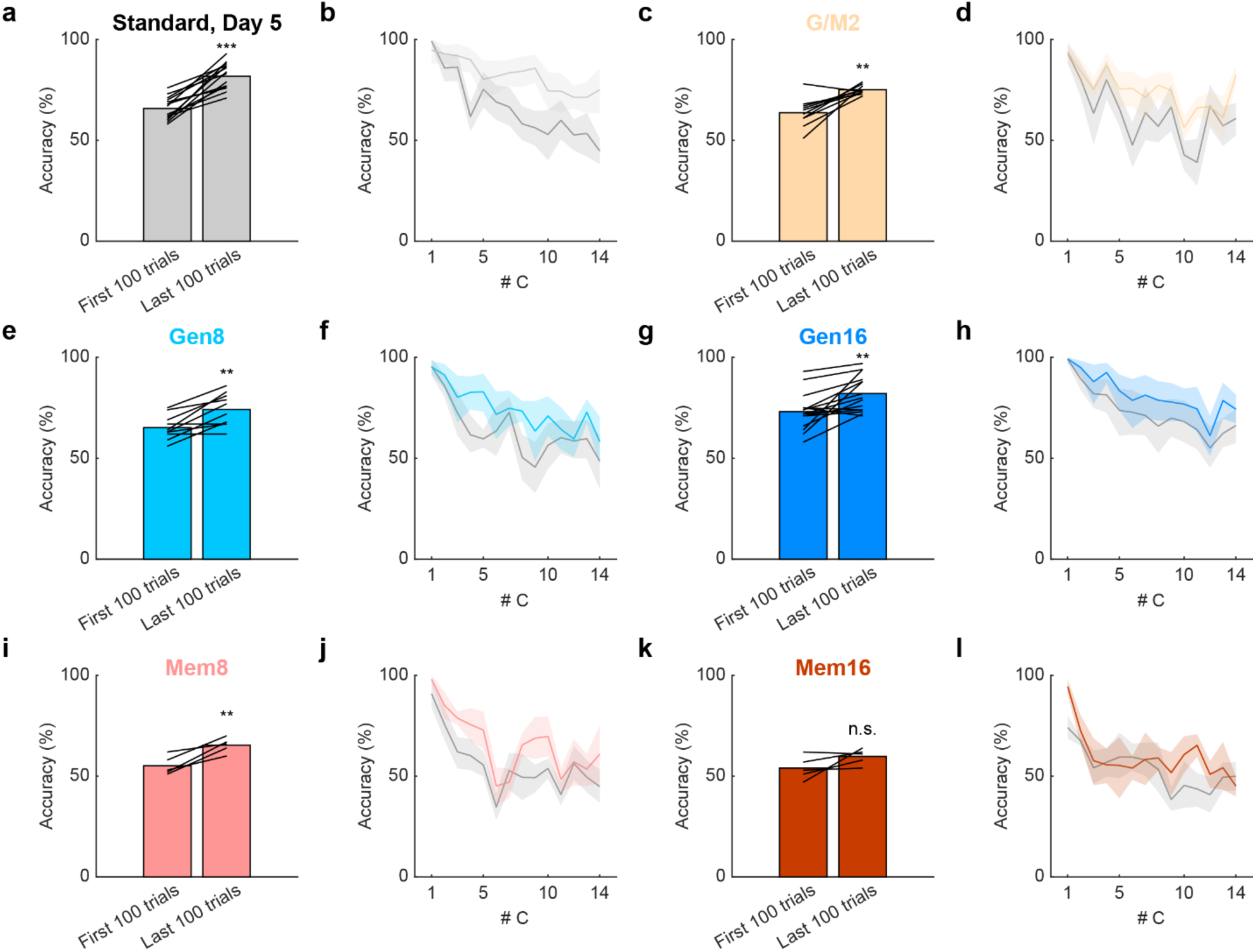
Learning within a session. (**a-b**) Within one session, animals improved their performance significantly. ***p<0.001, paired t-test. (**a**): Total accuracy. (**b**): Different accuracies to different #C. (**c-l**) Like (**a**), showing other groups of animals in different experimental conditions. Paired t-test.

**Figure S8:**
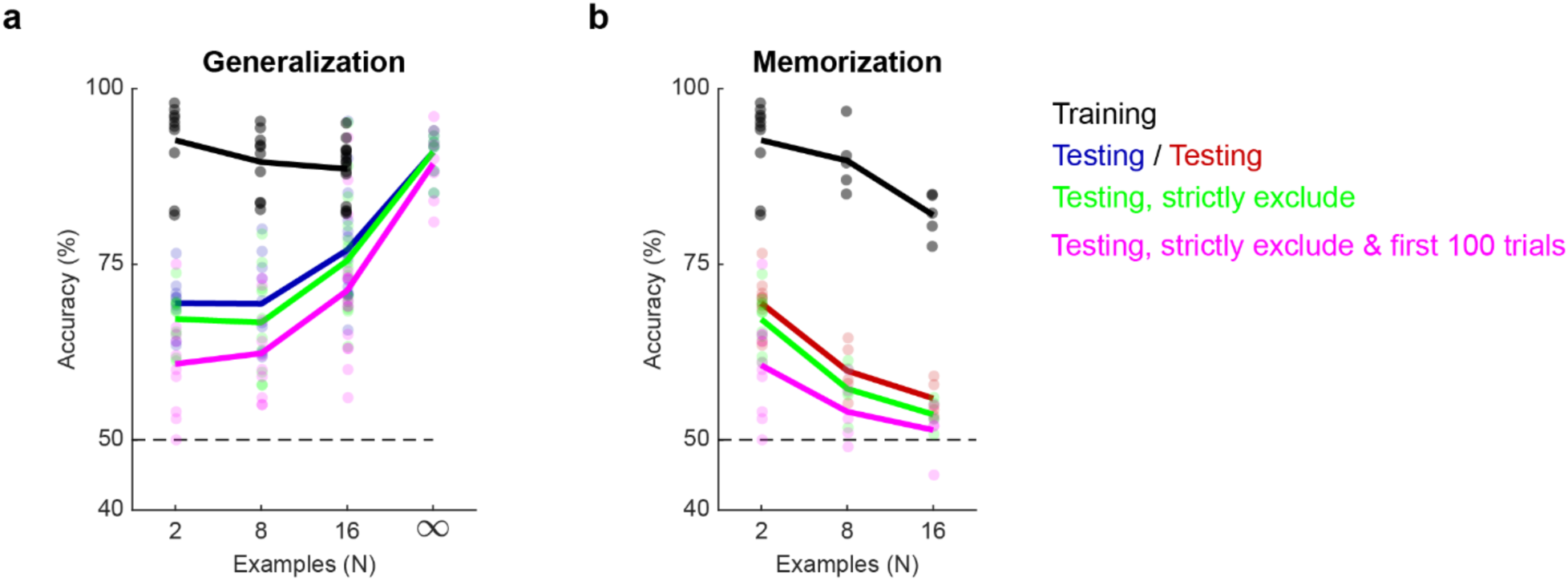
Strictly excluding training data from the testing session did not change the results. In Figure 2g and 2i, the test accuracy was calculated by all the trials in the session. With stricter criteria, the accuracies excluding trials with the same stimuli in the training set (green) or that plus restricting the calculation to the first 100 trials (magenta) are shown for (**a**): Generalization, and (**b**) Memorization experiments. With minor shifts, these treatments did not change our results or conclusions.

**Figure S9:**
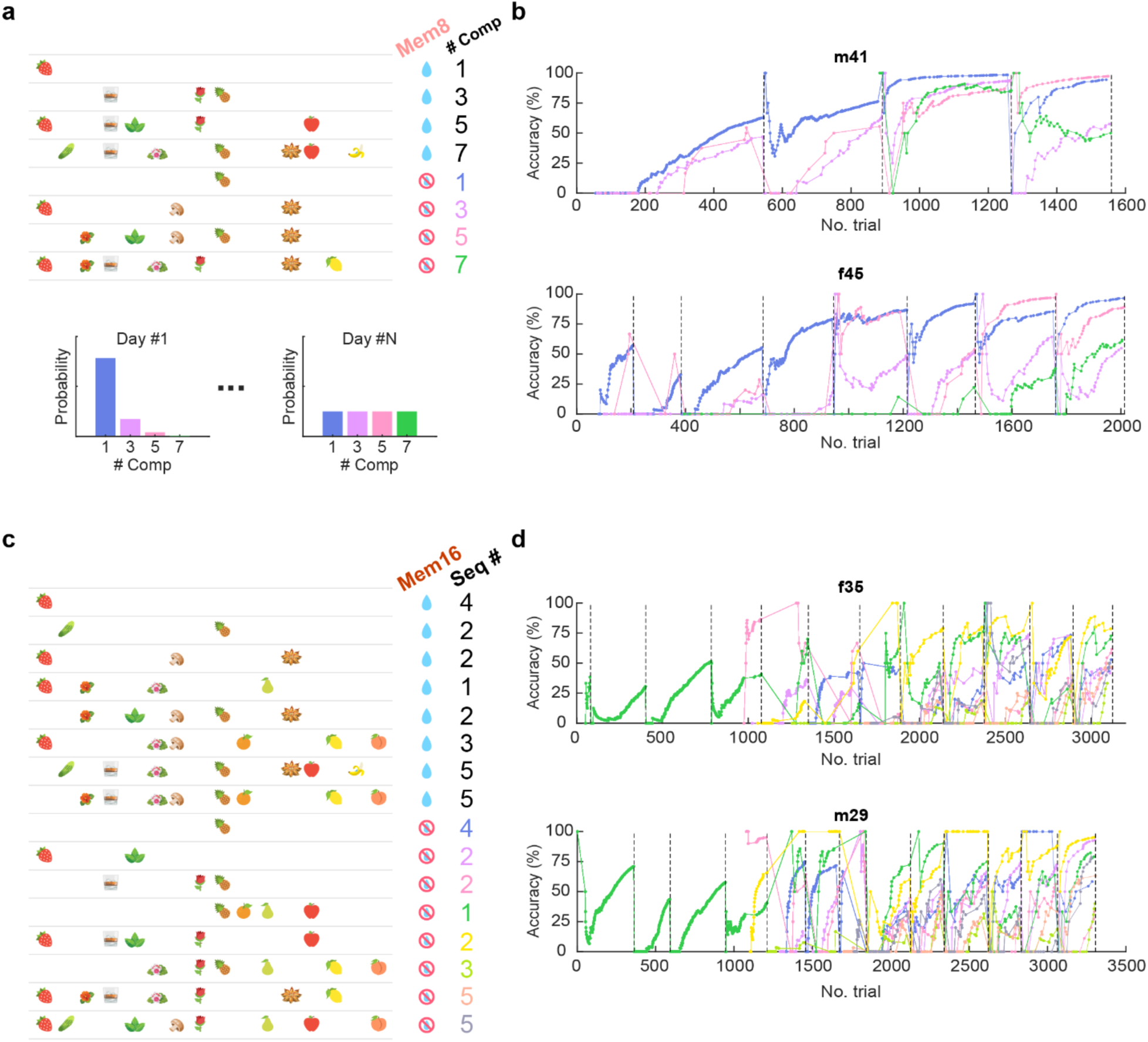
Training process of the memorization experiments. (**a**) Upper panel: the 8 mixtures and their labels in the Mem8 group. Mixtures are listed in increasing order of their number of components. The 4 NoGo mixtures are assigned with different colors, matched in the lower panel and (**b**). Lower panel: the probability of presenting different mixtures from early to later training days. (**b**) The complete training histories of two example animals in the Mem8 group, showing only the learning of NoGo odors. (**c**) The 16 mixtures and their labels in the Mem8 group. Mixtures are listed in increasing order of their number of components. Seq#: the sequence of training, from 1 to 5. The 8 NoGo mixtures are assigned with different colors, matched in (**d**). Unlike Mem8, a uniform distribution was used from the start, but restricted to the mixtures taught in sequence. (**d**) The complete training histories of two example animals in the Mem16 group, showing only the learning of NoGo odors.

**Figure S10:**
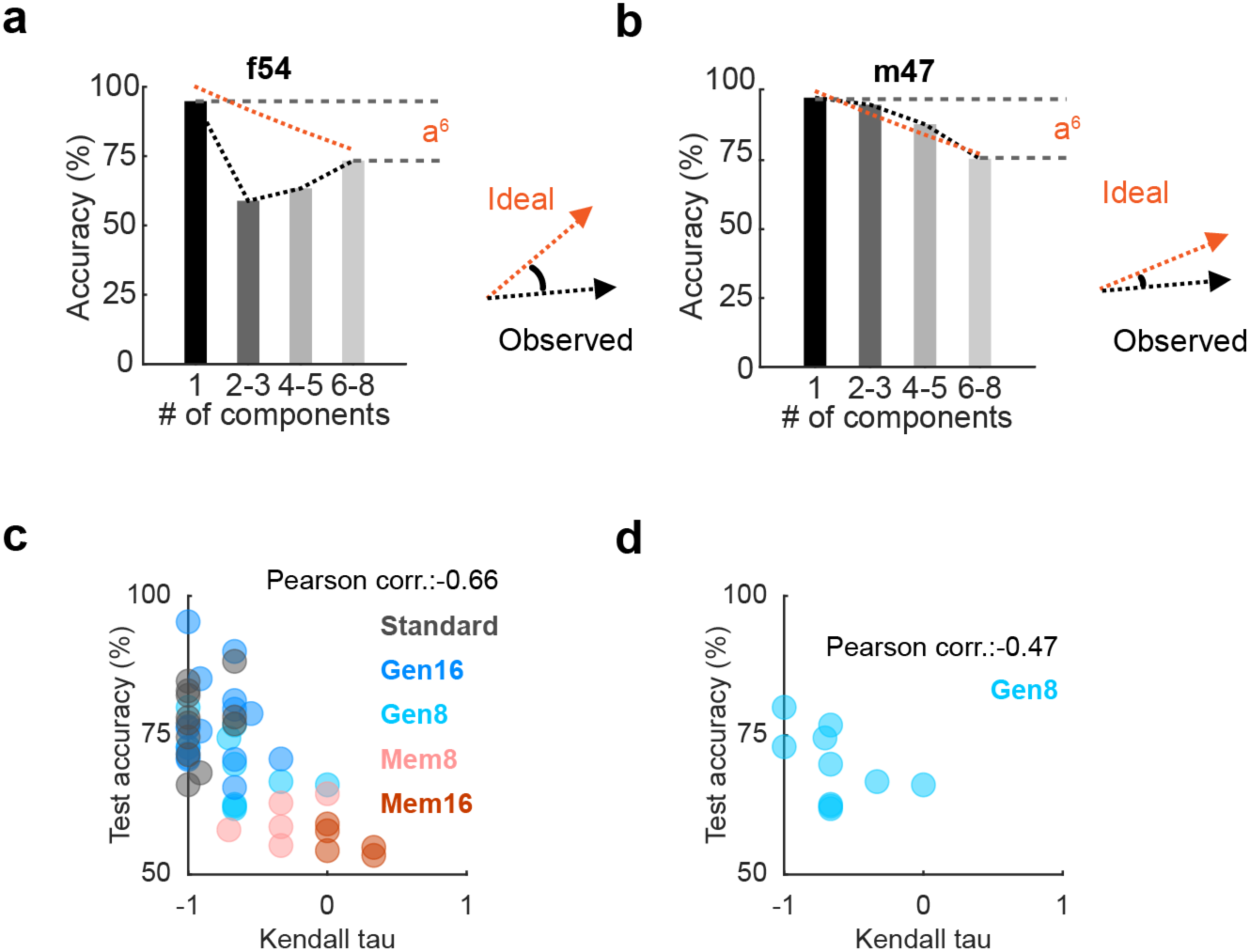
Two metrics to quantify order of learning in training. (**a-b**): Illustration of calculation of the metric θ, which reflects the angle between the ideal (red) and observed (black) 4-dimensional vector of accuracy to different #C mixtures: [C_1_,C_2-3_,C_4-5_,C_6-8_]. (**a**) Example mouse (“f54”) which did not match the ideal ordered sequence. (**b**) Example mouse (“m47”) which matched the ideal ordered sequence. (**c-d**) Using Kendall’s tau to measure the degree of rank disruption in the vector: [C_1_,C_2-3_,C_4-5_,C_6-8_] (compared to [1,2,3,4]), non-parametrically. (**c**) All groups of animals, N = 48 animals. (**d**) Gen8 group only, N = 10 animals.

**Figure S11:**
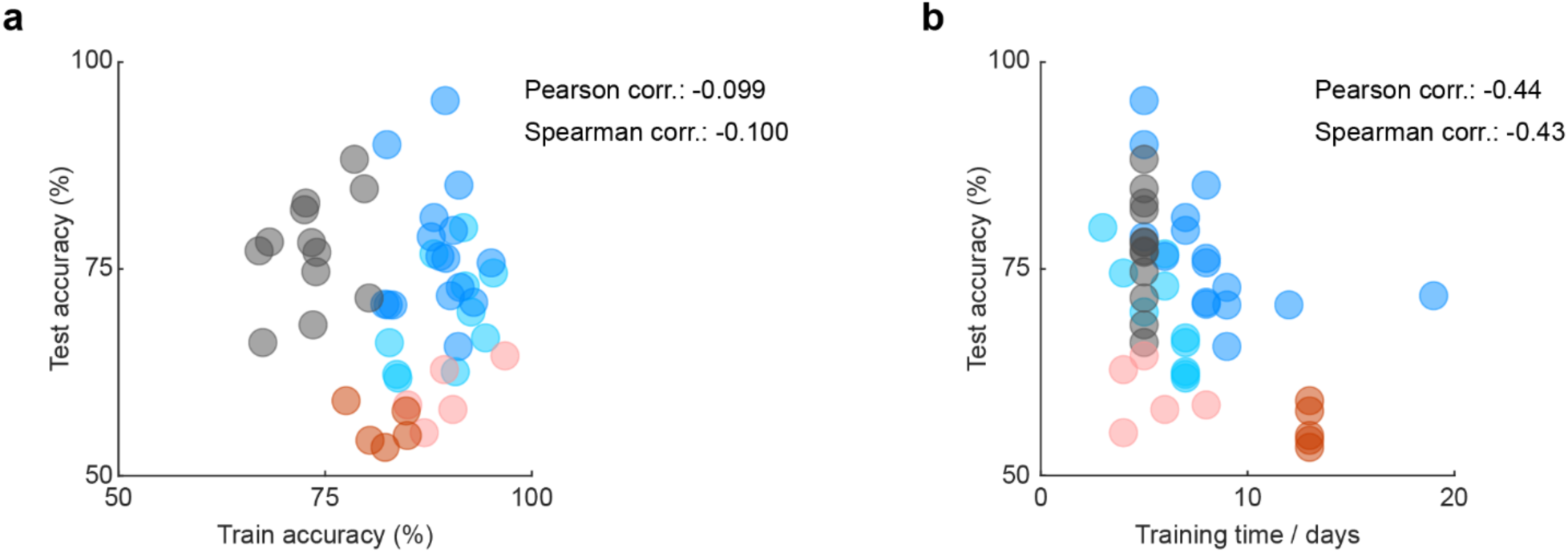
Training accuracy does not correlate with the testing accuracy, and display of the training days of all the groups. (**a**) Training accuracy does not correlate with the testing accuracy, showing that animals performing worse in the testing was not under trained in the training phase. (**b**) The training time differ across different groups due to different task demands, but across groups the training time still negatively correlates with the testing accuracy.

**Figure S12.**
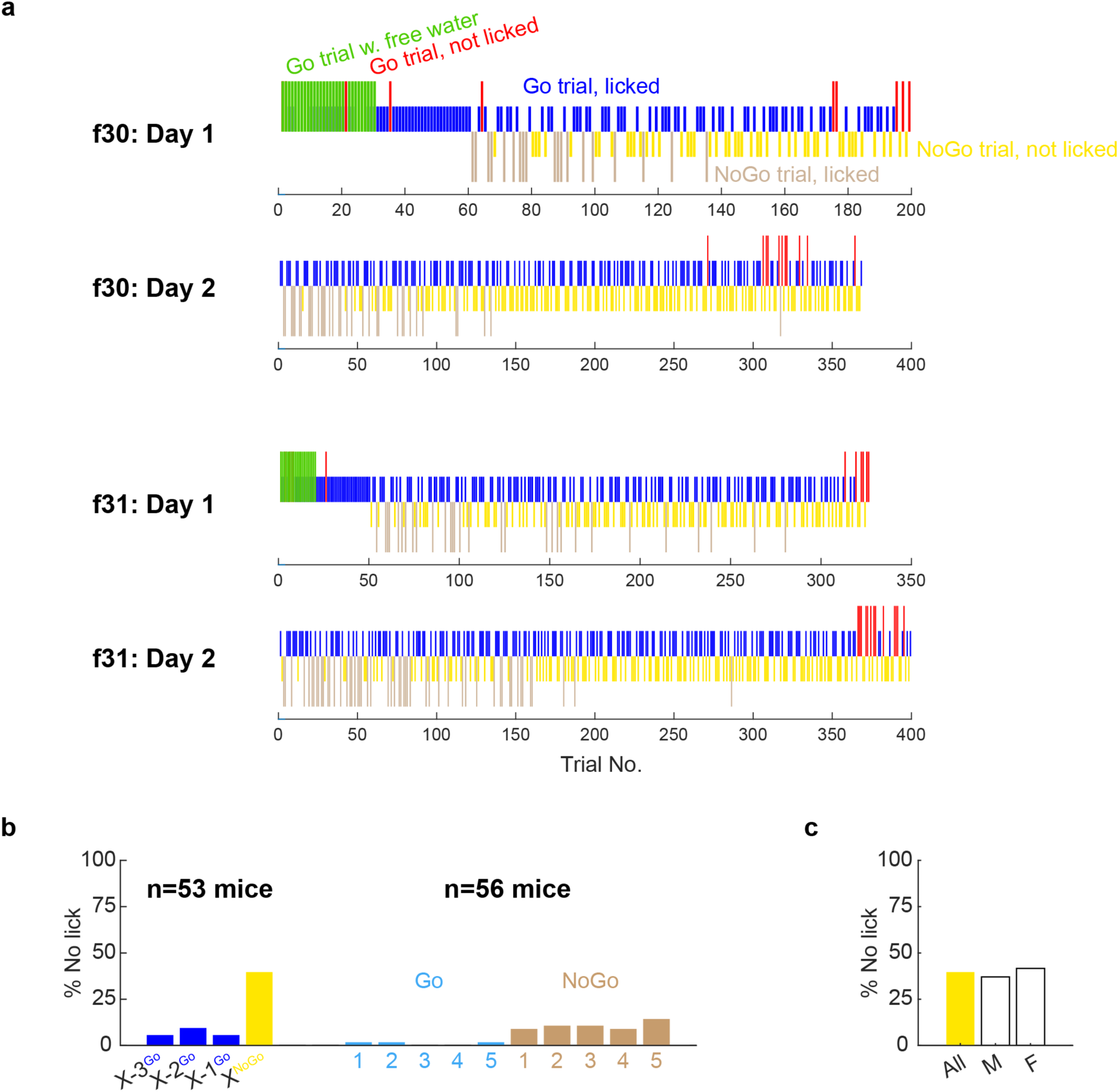
The novelty response was restricted to the beginning of training. (**a**) The complete training experience of the first two sessions (Day 1 and Day 2) of two example animals. Animals were first presented by consecutive Go trials (30-60 trials), starting with free water which was later removed. On the first encounter of the NoGo odor, some animals tried licking (f30) whereas others refrained from licking without any prior knowledge (f31). (**b**) Quantification of the percentage of not licking to the first 5 Go trials (light blue) and first 5 NoGo trials (light brown) on the next day. (**c**) No sex difference in the novelty response.

**Table S1.**
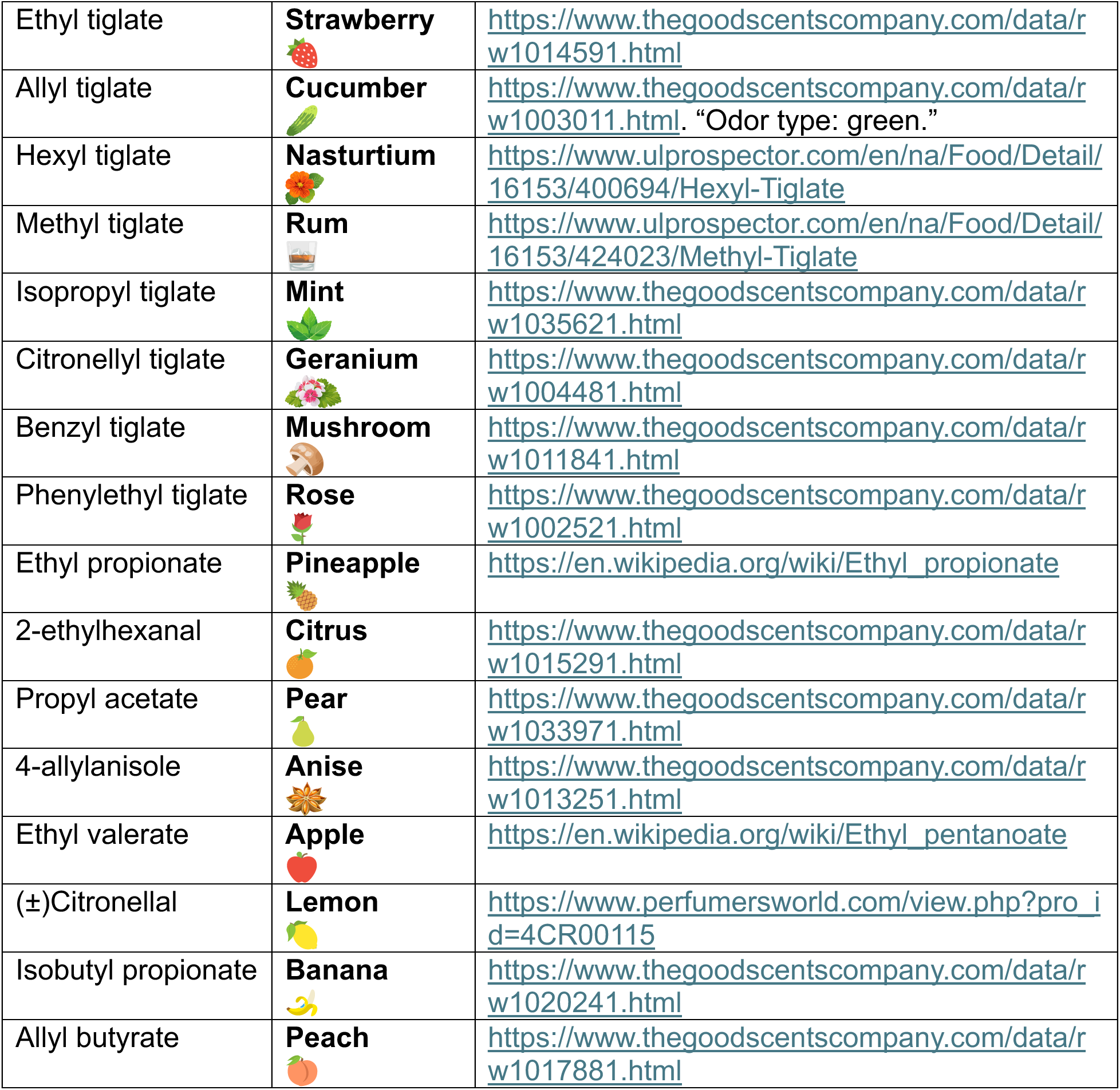
Description of smell and representative symbols of each odorant.

